# Amyloid fibrils in frontotemporal lobar degeneration with TDP-43 inclusions are composed of TMEM106B, rather than TDP-43

**DOI:** 10.1101/2022.01.31.478523

**Authors:** Yi Xiao Jiang, Qin Cao, Michael R. Sawaya, Romany Abskharon, Peng Ge, Michael DeTure, Dennis W. Dickson, Janine Y. Fu, Rachel R. Ogorzalek Loo, Joseph A. Loo, David S. Eisenberg

## Abstract

FTLD is the third most common neurodegenerative condition, following only Alzheimer’s and Parkinson’s diseases. FTLD typically presents in 45-64-year-olds with behavioral changes or progressive decline of language skills. The subtype FTLD-TDP is characterized by certain clinical symptoms and pathological neuronal inclusions detected by TDP-43 immunoreactivity. Here, we extracted amyloid fibrils from brains of four patients, representing four out of five FTLD-TDP subclasses and determined their near-atomic resolution structures by cryo-EM. Unexpectedly, all amyloid fibrils examined are composed of a 135-residue C-terminal fragment of TMEM106B, a lysosomal membrane protein previously implicated as a genetic risk factor for FTLD-TDP. In addition to TMEM106B fibrils, abundant non-fibrillar aggregated TDP-43 is present, as revealed by immunogold labeling. Our observations confirm that FTLD-TDP is an amyloid-involved disease and suggest that amyloid involvement in FTLD-TDP is of protein TMEM106B, rather than of TDP-43.

## Introduction

Pathological deposits of amyloid proteins are associated with over 50 systemic and neurodegenerative diseases^1,2^, including amyloid-β and tau in Alzheimer’s disease, α-synuclein in Parkinson’s disease, and hIAPP (or amylin) in type II diabetes^3^. Recent advances in cryogenic electron-microscopy (cryo-EM) techniques have enabled scientists to determine near-atomic resolution structures of amyloid fibrils extracted from patient tissues^4–9^. Structural information from *ex vivo* fibrils offers insight into the molecular signatures of disease, and can guide the design of therapeutics intended to prevent, delay or reverse the aggregation of proteins in amyloid disorders.

Frontotemporal lobar degeneration (FTLD) causes presenile dementia in ~81 out of 100,000 people between the ages of 45 to 64^10^. FTLD presents clinically as disorders of social behavior and language ability^11^. The major subtype of FTLD is characterized by neuronal inclusions containing TAR DNA-binding protein (TDP-43), termed FTLD-TDP, which accounts for ~50% of all FTLD cases^12^. FTLD-TDP is further classified into types A through E, according to morphology and neuroanatomical distribution of TDP-43 inclusions^13,14^; each type is associated with diverse clinical symptoms^15^. Although the defining neuropathological feature of FTLD-TDP is immunoreactivity for deposits of ubiquitinated, hyperphosphorylated TDP-43^16,17^, amyloid fibrils formed by TDP-43 or other proteins have not been reported in FTLD brains. Here, we extracted amyloid fibrils from four donors diagnosed with FTLD-TDP types A through D, and determined 12 near-atomic resolution structures by cryo-EM. By atomic model building, mass spectrometry, and western blot, we confirmed these fibrils are formed by transmembrane protein 106B (TMEM106B), a protein previously identified as a genetic risk factor of FTLD-TDP^18^. Our study reveals the amyloidogenic nature of TMEM106B in FTLD-TDP, and focuses attention on a protein previously not associated with amyloid conditions^19–21^.

## Results

### Fibril extraction from FTLD-TDP patients and cryo-EM study

The Mayo Clinic Brain Bank contributed frozen brain tissues of FTLD-TDP patients (40 donors) and age-matched, non-FTLD-TDP controls (8 donors, Fig. S1a and Table S1). We inspected the detergent-insoluble fractions of these samples by negative stain transmission election microscopy. In most donors with FTLD-TDP (38 out of 40 donors), we observed amyloid fibrils as well as non-fibrillar aggregates (donors 1-4 in Fig. 1a, and 34 other donors in Fig. S2). In contrast, no fibrils were observed in all 8 non-FTLD-TDP donors, (donor 5 in Fig. 1a, and 7 other donors in Fig. S2). We then selected four donors neuropathologically confirmed to be FTLD-TDP types A, B, C, and D (donors 1-4, Fig. S3 and Table 1) for further study. We collected cryo-EM data on samples from each of the four donors and determined amyloid fibril structures (Fig. 2, Fig. S4–6, Table 2 and Table S2). Three fibril polymorphs, PM1, PM2, and PM3, were identified in each cryo-EM dataset (Fig. S4a-c), with similar distributions in all four FTLD-TDP donors (Fig. 2 and Fig. S4d).

**Fig. 1.**
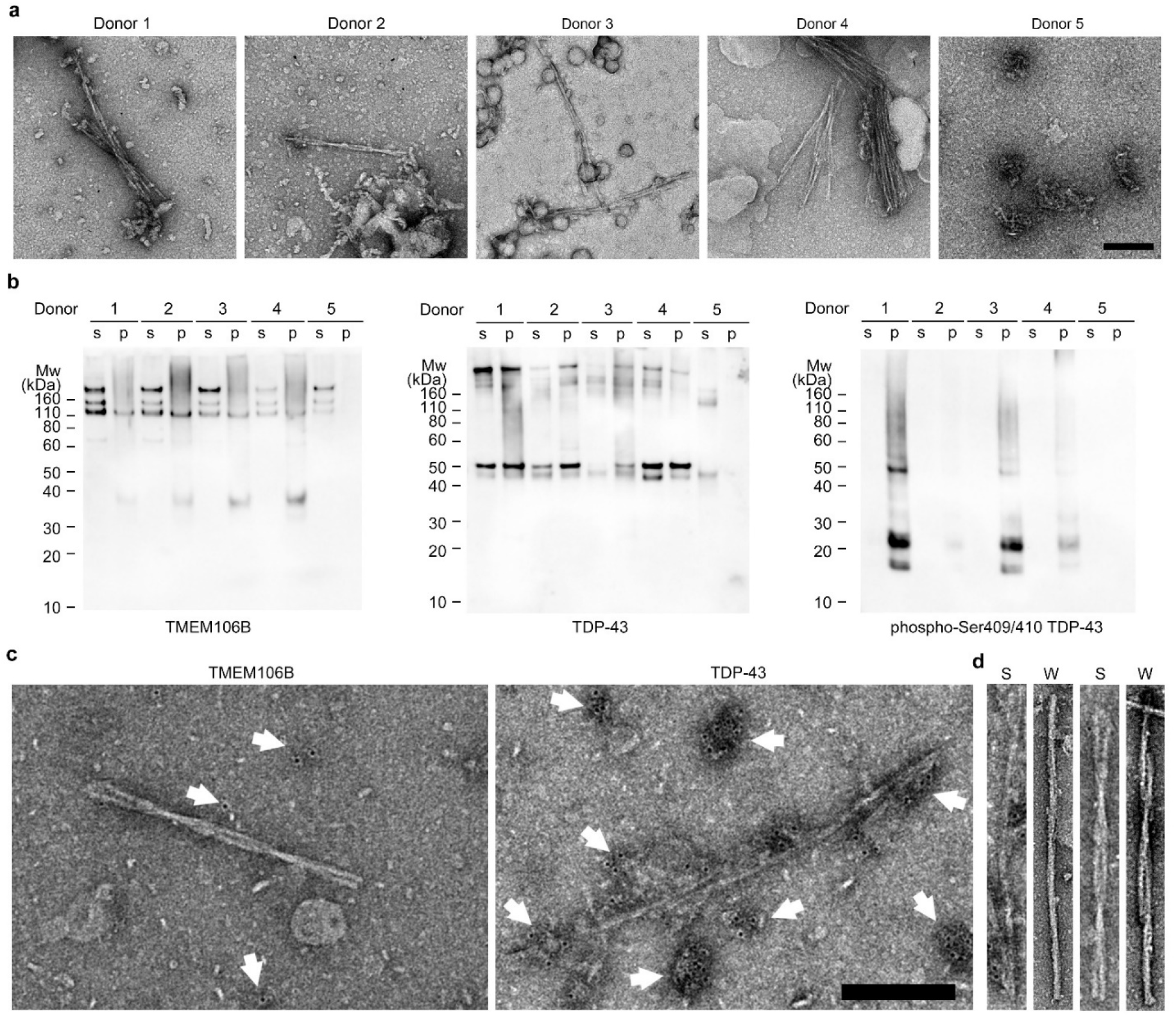
Characterization of TMEM106B and TDP-43 in FTLD-TDP brain extracts. **a**, Negative stain TEM images of sarkosyl-insoluble fractions from FTLD-TDP donors 1 to 4 and a non-FTLD-TDP donor 5. Scale bar 200 nm. **b**, Western blots of sarkosyl-soluble (s) and sarkosyl-insoluble (p) fractions from donors 1 to 5 probed with TMEM106B (left), TDP-43 (middle), or phospho-Ser409/410 TDP-43 (pTDP-43, right) antibody. **c**, Representative negative stain TEM images of the sarkosyl-insoluble fraction from FTLD-TDP donor 1 showing immunogold labeling using TMEM106B (left) and TDP-43 (right) antibodies. White arrows label immunogold beads, showing low density, likely nonspecific binding of TMEM106B antibody (left) and high density, non-fibrillar TDP-43 aggregates (right). Scale bar 200 nm. **d,** Comparison of fibrils extracted from FTLD-TDP donor 1 using sarkosyl-based protocol (S, adopted from c) and water-based protocol (W) by negative stain TEM. PM1(left) and PM2 or 3 (right, PM2 and PM3 are difficult to distinguish by negative stain TEM). Fibrils extracted from both protocols exhibit similar morphology.

**Fig. 2.**
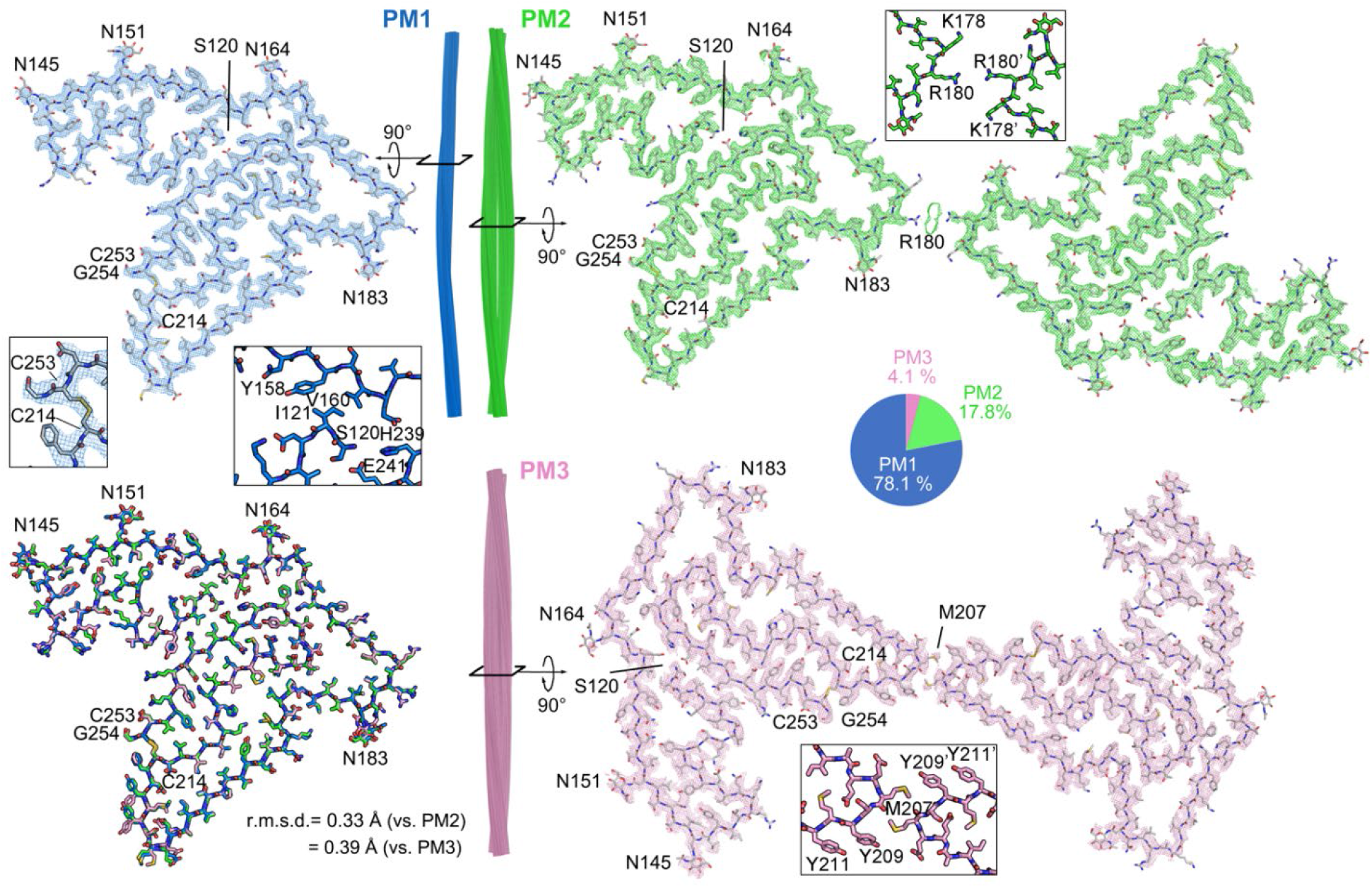
Cryo-EM structures of TMEM106B fibrils from FTLD-TDP donor 1. Cryo-EM maps and atomic models of one cross-sectional layer of PM1 (blue, top left), PM2 (green, top right), and PM3 (pink, bottom right). Side view of fibril reconstructions of PM1, PM2 and PM3 (middle). Superimposition of a single chain from PM1, PM2 and PM3 (bottom left). Enlarged views of the C214-C253 disulfide bond, N-terminal Ser120 in PM1, and dimer interfaces of PM2 and PM3 are shown as inserts. Distribution of PM1, PM2, and PM3 polymorphs in the donor 1 dataset is shown as a pie chart.

**Table 1.** Information of FTLD-TDP and non-FTLD-TDP donors.

| Donor ID | Gender | Age | Braak | PathDx (type) | FHx | Brain region | Distribution (%) |  |  | TMEM106B genotype |
| --- | --- | --- | --- | --- | --- | --- | --- | --- | --- | --- |
|  |  |  |  |  |  |  | PM1 | PM2 | PM3 |  |
| Donor 1 | Male | 65 | 2 | FTLN-TDP (C) | False | MF | 78.1 | 17.8 | 4.1 | Heterozygous T185S |
| Donor 2 | Female | 76 | 3 | FTLN-TDP (B) | False | MF | 84.9 | 10 | 5.1 | Heterozygous T185S |
| Donor 3 | Male | 86 | 0 | FTLN-TDP (A) | False | MF | 85.3 | 12.8 | 1.8 | Wildtype |
| Donor 4 | Female | 64 | 3 | FTLN-TDP (D) | True | MF | 85.3 | 10.7 | 4 | Heterozygous T185S |
| Donor 5 | Male | 59 | 0 | Normal | False | MF | n.a. | n.a. | n.a. | n.a. |

PathDx: pathological diagnosis; FHx: family history; MF: medial frontal gyrus; n.a.: not available.

**Table 2.** Cryo-EM data collection, refinement and validation statistics of FTLD-TDP donor 1.

|  | PM1<br>(EMD-24953,<br>PDB 7SAQ) | PM2<br>(EMD-24954,<br>PDB 7SAR) | PM3<br>(EMD-24955,<br>PDB 7SAS) |
| --- | --- | --- | --- |
| **Data collection and processing** |  |  |  |
| Magnification | ×81,000 | ×81,000 | ×81,000 |
| Voltage (kV) | 300 | 300 | 300 |
| Electron exposure (e<sup>-</sup>/Å<sup>2</sup>) | 36 | 36 | 36 |
| Defocus range (μm) | 0.5-5.0 | 0.5-5.0 | 0.5-5.0 |
| Pixel size (Å) | 1.1 | 1.1 | 1.1 |
| Symmetry imposed | C<sub>1</sub> | C<sub>2</sub> | C<sub>1</sub> |
| Helical rise (Å) | 4.91 | 4.905 | 2.452 |
| Helical twist (°) | -0.42 | 179.59 | 179.79 |
| Initial particle images (no.) | 883,793 | 883,793 | 883,793 |
| Final particle images (no.) | 60,806 | 16,883 | 9,203 |
| Map resolution (Å) | 2.9 | 3.2 | 3.7 |
| FSC threshold | 0.143 | 0.143 | 0.143 |
| Map resolution range (Å) | 200-2.9 | 200-3.2 | 200-3.7 |
| **Refinement** |  |  |  |
| Initial model used (PDB code) | De novo | 7SAQ | 7SAQ |
| Model resolution (Å) | 3.1 | 3.4 | 4.3 |
| FSC threshold | 0.5 | 0.5 | 0.5 |
| Model resolution range (Å) | 200-3.1 | 200-3.4 | 200-4.3 |
| Map sharpening *B* factor (Å<sup>2</sup>) | 124 | 93 | 98 |
| Model composition |  |  |  |
| Nonhydrogen atoms | 5,710 | 11,420 | 11,420 |
| Protein residues | 675 | 1,350 | 1,350 |
| Ligands | 20 | 40 | 40 |
| *B* factors (Å<sup>2</sup>) |  |  |  |
| Protein | 7.10 | 6.57 | 29.53 |
| Ligand | 11.64 | 10.61 | 52.32 |
| R.m.s. deviations |  |  |  |
| Bond lengths (Å) | 0.005 | 0.004 | 0.004 |
| Bond angles (°) | 0.793 | 0.732 | 0.818 |
| **Validation** |  |  |  |
| MolProbity score | 2.21 | 2.27 | 2.37 |
| Clashscore | 11.87 | 15.13 | 18.56 |
| Poor rotamers (%) | 0 | 0 | 0 |
| Ramachandran plot |  |  |  |
| Favored (%) | 87.22 | 88.64 | 87.97 |
| Allowed (%) | 12.78 | 11.36 | 12.03 |
| Disallowed (%) | 0 | 0 | 0 |

### Atomic model building and identification of the fibril-forming protein

We used our highest resolution cryo-EM map (2.9 Å, from donor 1, PM1) to begin model building. We presumed the fibrils were composed of TDP-43; however, extensive incompatibility between the side chain density in the map and the sequence of TDP-43 invalidated this presumption. To identify which protein actually composes the fibrils, we constructed query sequences by choosing amino acids that best fit the PM1 density without regard to their similarity to known protein sequences. We built two models having opposite directionality, thereby obtaining two query sequences (Fig. S7, see Methods). A search for human proteins similar to these two queries revealed only one hit: TMEM106B (residues 121-254), a lysosomal transmembrane protein.

Several observations lend confidence that TMEM106B is indeed the protein building block of these fibrils: i) the sequence of TMEM106B fits the map well leaving no unexplained density (Fig. 2); ii) density protruding from Asn145, Asn151, Asn164, and Asn183 (Fig. S8a&b) accounts for the known N-linked glycosylation sites of TMEM106B (Fig. 3a)^22^; iii) the only density that connects two side chains is accounted by a disulfide bond between Cys214 and Cys253 (Fig. 2); iv) a previous study revealed that genetic variants of TMEM106B are associated with FTLD-TDP^18^. No other protein could explain these features so completely. The models for PM2 and PM3 of donor 1 were built by rigid body fits of the PM1 model into the density maps, with minor adjustments and refinement applied subsequently.

**Fig. 3.**
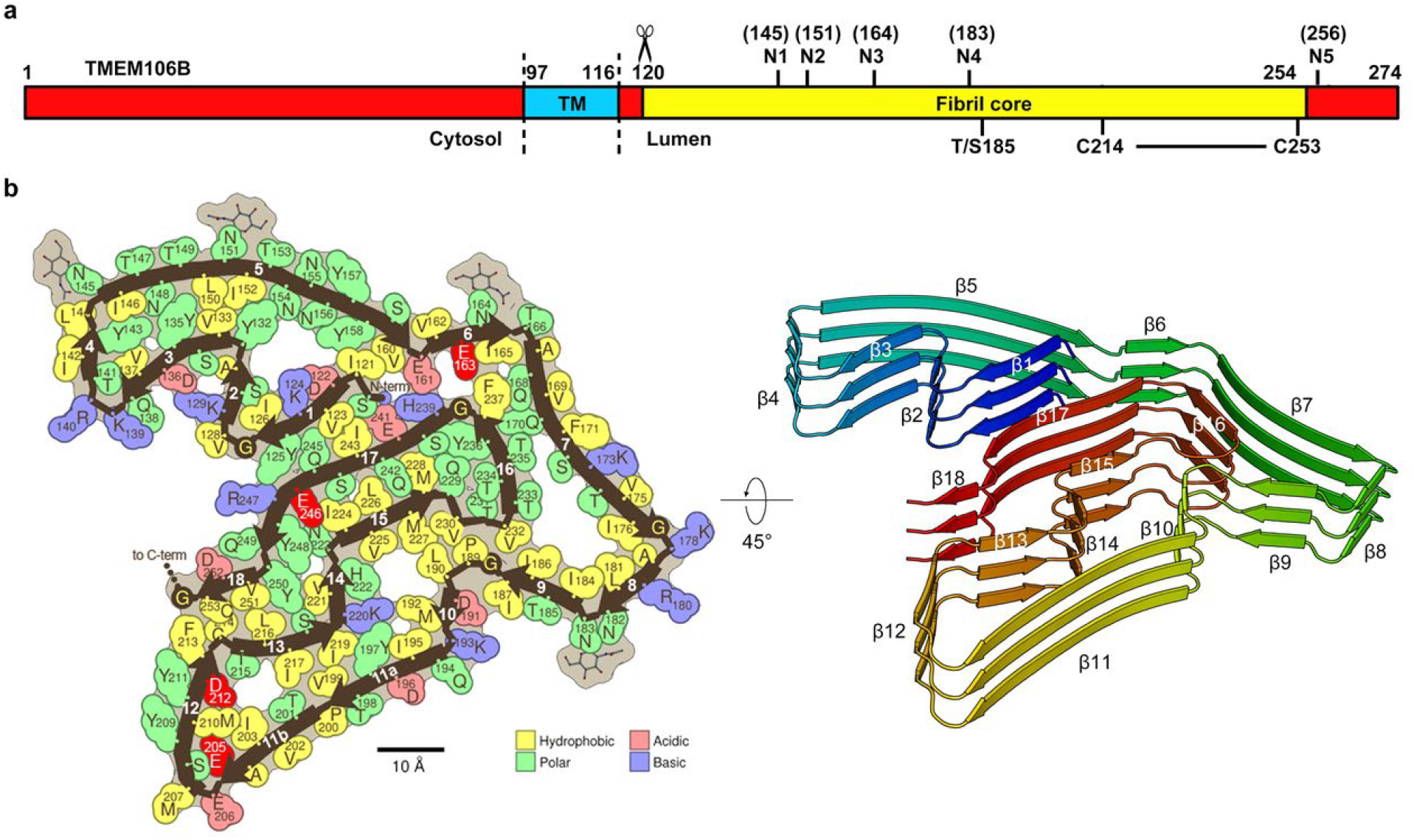
The conserved protofilament golf-course-like fold of TMEM106B fibrils. **a**, Schematic of TMEM106B with the residue ranges of the cytosolic, transmembrane (TM) and lumenal domains annotated, as well as the fibril core of the structures observed here. Labelled are the five known glycosylation sites (N1-N5), the heterozygous Thr185Ser variant, and the C214-C253 disulfide bond. **b**, Space-filling representation (left) and cartoon model (right) of the conserved protofilament core represented by PM1 of FTLD-TDP donor 1. Uncomplemented acidic residues buried inside the fibril core are highlighted in bright red. N-term, N terminus; C-term, C terminus.

Cryo-EM maps of all three polymorphs make it clear that the fibrils are composed of a proteolytic fragment, rather than the full-length TMEM106B. The N-terminus of the core, Ser120, is buried by residues Tyr158, Val160, His239 and Glu241, leaving no space for residues preceding Ser120 (Fig. 2 and Fig. 3b). Consequently, we conclude that proteolytic cleavage between residues Arg119 and Ser120 must precede fibril formation (Fig. 3a, Fig. S9a, see Discussion). In contrast, the C-terminus of the core, Gly254, is solvent-exposed, allowing the possibility that the remaining C-terminal 20 residues are attached but disordered. Notably, the range of the ordered core (residues 120-254) coincides closely with the definition of the lumenal domain of TMEM106B (Fig. 3a), which is known to be released from the lysosomal membrane by proteolytic cleavage at an undetermined location within the vicinity of the site we infer from our structure, Ser120^23^.

### Polymorphic fibrils extracted from donor 1 share a stable, conserved golf course-like fold

Protofilaments of all three polymorphs of donor 1 exhibit the same fold (Fig. 2, bottom left) despite their different states of bundling. A single protofilament constitutes PM1, while a pair of protofilaments constitute PM2 and PM3 and intertwine with distinct symmetries: C_2_ in PM2 and pseudo-2_1_ in PM3. Moreover, the protofilament interfaces of PM2 and PM3 differ. In PM2, side chains of Arg180 point toward residual density at the center of the interface, whereas PM3 features a hydrophobic interaction between pairs of Met207 and Tyr209 residues (Fig. 2 and Fig. S10). The only notable structural differences between the folds in all three polymorphs are confined to the PM3 protofilament interface; the rotamers of Glu206, Met207, Tyr209, and Tyr211 in PM3 vary from those in PM1 and PM2 (Fig. 2 and Fig. S10c).

We liken the conserved fold of TMEM106B to a golf course because it contains 18 β-strands (Fig. 3b and Fig. S9a), corresponding to the conventional number of fairways in a golf course. These 18 β-strands vary in length (from 3 to 15 residues) and curvature as do the fairways in a golf course. Moreover, the N-and C-terminal strands are near each other, just as the 1^st^ and 18^th^ holes in a golf course adjoin near the clubhouse.

Importantly, although 18 holes are typical of a golf course, 18 β-strands exceed the number typical of amyloid fibrils^24^; this abundance of β-strands contributes to an unusually large and stable fibril core. Indeed, these 18 β-strands mate together in pairs, forming multiple steric zippers that strongly stabilize the fibril (Fig. S9b). Compared to the average of 63 amyloid fibril structures known in 2021, the golf course fold of TMEM106B ranks nearly three times more stable by solvation energy estimates^24^ (−62 vs. −22 kcal/mol/chain). These crude estimates of energetic stability may explain the fibrils’ resistance to 2% sarkosyl and 1% SDS during extraction. Such stability suggests that formation of TMEM106B fibrils in the brain is irreversible, which is consistent with their possible pathogenic roles in FTLD-TDP.

Clues to the location of origin of TMEM106B fibrils are revealed by our observation of four uncomplemented acid side chains face inward in the fibril core structure: Glu163, Glu205, Asp212, and Glu246 (Fig. 3b). Such uncomplemented, buried negative charges are rare among pathogenic amyloids as electrostatic repulsion between closely spaced molecules with the identical charge would weaken the fibril. However, considering TMEM106B is a lysosomal protein and this is a lumenal domain, the pH of its environment might be low enough to protonate and neutralize some inward facing acids. Neutralization of these negative charges may help to overcome the barrier to fibril nucleation.

### FTLD-TDP donors 1-4 share a conserved fold despite different subtypes and genotypes

Our study of four FTLD-TDP patients contributes to mounting evidence that protein conformation is conserved among patients diagnosed with the same amyloid disease. In fact, it has been proposed that protein conformation could form the basis for classifying disease^25^. Accordingly, we found the golf course fold of TMEM106B is conserved among all four FTLD-TDP donors and constitutes the basis of all three polymorphs. However, we do observe a small but significant shift in the protofilament-protofilament interface of PM2 among donors. The interface from donors 1 and 4 (types C and D) is centered on Lys178, whereas the interfaces from donors 2 and 3 (types B and A) is centered on Arg180 (Fig. S6 and Fig. S10a&b). It is unclear whether these structural differences are linked to the different FTLD-TDP subtypes or other patient attributes, such as age. PM2 accounts for ~10% of TMEM106B fibrils in all four donors

Genotyping suggests that donor 3 contains wildtype TMEM106B, whereas donors 1, 2, and 4 harbor heterozygous Thr185Ser variants in their TMEM106B gene (Table 1). Mass spectrometry identified a peptide with Thr185Ser variant in donor 1 (Fig. S11 and Table S3). Thr185Ser variant was detected in a genome-wide association study for FTLD-TDP patients^18^, and a follow-up study suggested the Thr185Ser variant has protective effect against FTLD-TDP^26^. The Ser185 isoform is associated with lower protein expression and is more rapidly degraded than the wildtype Thr185 isoform, which may contribute to its protective mechanism^27^. The observation that all four donors possess fibrils with the same fold suggests that this fold is compatible with wildtype TMEM106B protein; however, we cannot confirm whether the heterozygous donor’s fibrils contain only wildtype, only variant, or a mixture of both. Our maps confirm that residue 185 is within the protofilament core (Fig. 2a and Fig. S8c), but even the highest resolution cryo-EM map (donor 1 PM1) is insufficient to distinguish among Thr, Ser, or an average of both sidechains (Fig. S8c). Since the Thr/Ser185 side chain faces the solvent with no contributing interactions in the fibril structure, we believe the fibril fold observed here can accommodate both wildtype and Thr185Ser TMEM106B proteins.

### Immunostaining and mass spectrometry verify TMEM106B and TDP-43 are present in FTLD-TDP patient extracts

To verify that the protein in FTLD-TDP fibrils is indeed TMEM106B, we performed western blotting of patient extracts with a C-terminal TMEM106B antibody. We observed a ~35 kDa TMEM106B-positive band (along with other high molecular weight bands, see Methods) in sarkosyl-insoluble fractions of most of the FTLD-TDP donors examined but none of the non-FTLD-TDP controls (Fig. 1b and Fig. S1b), consistent with our observations by EM that TMEM106B fibrils are present only in FTLD-TDP patients (Fig. 1a and Fig. S2). These results support the hypothesis that the aggregation of TMEM106B is associated with FTLD-TDP. The ~35kDa band could be the fibril-forming cleavage product (TMEM106B 120-274), a 155-residue peptide with five glycans. To further confirm the presence of TMEM106B, we performed mass spectrometry on gel slices around the ~35kDa region and detected peptide fragments from only the lumenal domain (Fig. S11 and Table S3), consistent with our cryo-EM maps.

We attempted immunogold labeling of patient-extracted fibrils using three different TMEM106B antibodies (see Methods), and none labeled the fibrils (Fig. 1c, left). This was anticipated, since the epitopes of the antibodies are either cleaved before fibril formation or buried inside the fibril core. It may also explain why TMEM106B aggregation has not been detected by immunohistochemical staining of patient brains. Future work is needed to develop antibodies that probe TMEM106B fibrils, either by binding to the fibril structure, or by recognizing residues 255-274 of TMEM106B, which are not included in the fibril core and may form an accessible “fuzzy coat”.

To verify the presence of TDP-43 aggregates, we also probed patient extracts with TDP-43 antibodies. Immunoblotting revealed that the FTLD-TDP-associated, phosphorylated form of TDP-43^28^ is present with TMEM106B in the sarkosyl-insoluble fractions of FTLD-TDP donors but not the non-FTLD-TDP control (Fig. 1b). These results are consistent with a previous study^29^ and suggest TDP-43 forms aggregates in FTLD-TDP patients. Immunogold labeling of sarkosyl-insoluble fractions with the TDP-43 antibody revealed non-fibrillar aggregates were present and identified them as TDP-43(Fig. 1c, right, see Discussion). Fibrils were also detected but were not labeled; presumably composed of TMEM106B.

## Discussion

A decade ago, TMEM106B was identified as a genetic risk factor for FTLD-TDP in a genome-wide association study; ever since, its role in the disease has remained elusive. Our finding of TMEM106B amyloid fibrils in *ex vivo* patient brains gives us a firm, new grasp on this association. Indeed, TMEM106B amyloid appears to be a signature of FTLD-TDP as evidenced by its presence in four different FTLD-TDP types, both with and without the Thr185Ser variant, and in both familial and sporadic cases (Table 1). Each of the three polymorphic structures we observe from each patient reveals TMEM106B folded in the same golf-course-like pattern. Our discovery directs new focus on TMEM106B and the role that its aggregation may have on the aggregation of TDP-43, previously recognized as “the major disease protein”^30^. Are TMEM106B fibrils pathogenic? If so, does pathogenicity arise from a gain or loss-of-function? Or, are TMEM106B fibrils benign by-products downstream of an undiscovered, primary pathological pathway?

Our structures reveal that proteolytic cleavage happens prior and is essential to form the core of TMEM106B fibrils (Fig. 2 and Fig. 3a), if so protease inhibition might offer a strategy for FTLD-TDP intervention. TMEM106B undergoes lumenal domain shedding, in which the amyloid core-forming C-terminal fragment is released within the lumen of the lysosome by a resident protease; the cleavage site and the identity of the protease is still unknown^23^. Our structural evidence that cleavage occurs between Arg119 and Ser120 (Fig. S12) is consistent with the propensity of many known proteases to cleave between arginine and serine residues. The specificity of the TMEM106B cleavage site required to enable fibril growth would seem to be even greater than that of amyloid precursor protein^32^, another single-pass transmembrane protein whose proteolytic cleavage by secretases is infamously involved in the growth of amyloid fibrils in Alzheimer’s disease. In the case of β-amyloid, cleavage at various sites leads to amyloid formation, whereas in TMEM106B, burial of its cleaved terminus in the fibril core constrains amyloid-competent fragments to have been cleaved between Arg119 and Ser120.

Evidence of a ligand binding site in the TMEM106B PM2 polymorph identifies another potential factor affecting fibril formation. The interface between PM2 protofilaments features positively-charged residues (Lys178 or Arg180) that point to an unmodeled residual density (Fig. S10b). This positively-charged environment suggests that the ligand bound here is negatively-charged. Our observation that this ligand accommodates two slightly different protofilament arrangements of PM2 (Fig. S10a) is consistent with the small size of the interface and the dominance of electrostatics in its stabilization. Interestingly, residual densities are also found in similar positions in PM1 and PM3 (Fig. S8a, indicated by red arrows), and may correspond to the same ligand. Unfortunately, the resolution of the residual densities does not allow us to identify the ligand. We cannot rule out the possibility that the density corresponds to sarkosyl, a negatively-charged compound at pH 7.5, that may have been incorporated during fibril extraction. If fibril formation is sensitive to the presence of this ligand, then ligand withdrawal may inhibit their formation. Ligand binding sites have similarly been discovered in pathogenic fibrils of tau^9^ and α-synuclein^8^.

Our experiments show that aggregates of TDP-43 in FTLD-TDP are amorphous, not amyloid-like. FTLD-TDP has been defined by inclusions of TDP-43^12^. However, sarkosyl-insoluble TDP-43 extracted from FTLD-TDP patients exhibits non-fibrillar morphology rather than fibrillary morphology, shown both in our work (Fig. 1) and a previous study^29^. Our cryo-EM structures suggest that the amyloid fibrils we imaged contain only TMEM106B; however, we cannot exclude the possibility that TDP-43 forms fibrils in FTLD-TDP patients, and our extraction method failed to capture TDP-43 fibrils. We believe this is unlikely because we observed no fibrils in other fractions throughout our fibril extraction protocol by EM, and the sarkosyl and SDS treatment should not dissolve pathogenic irreversible fibrils. From our western blot and immunogold labeling experiments, we found TDP-43 aggregates can be co-extracted with TMEM106B fibrils from FTLD-TDP patients. It is so far unclear whether TDP-43 aggregates and TMEM106B fibrils co-localize in patients’ brains, and whether a cross-seeding mechanism is involved in FTLD-TDP pathogenesis.

### Conclusion

Our structures of patient-derived TMEM106B fibrils reveal a previously unrecognized process: protein cleavage followed by amyloid formation of the C-terminal fragment of TMEM106B. This finding may refocus the pathogenic studies of FTLD-TDP and perhaps other neurodegenerative diseases to include TMEM106B.

## Acknowledgements

We thank H. Zhou for the use of Electron Imaging Center for Nanomachines (EICN) facilities and Drs. Sjors Scheres and Michel Goedert for discussion of TMEM106B antibodies. We acknowledge the use of EICN instruments supported by the NIH (1S10RR23057 and IS10OD018111), NSF (DBI-1338135) and California NanoSystems Institute at UCLA. We thank National Facility for Translational Medicine (Shanghai) for support. The authors acknowledge NIH AG 054022, NIH AG061847, and DOE DE-FC02-02ER63421 for support.

## Author Contributions

Y.J. and Q.C. extracted fibrils from FTLD-TDP patients, prepared cryo-EM grids, collected and processed cryo-EM data. Y.J., Q.C., and M.R.S. built the atomic models. P.G. assisted in cryo-EM data collection. Y.J. and R.A. performed western blotting and immunolabeling assays. Y.J. performed patient genotyping. J.Y.F, R.R.O.L., and J.A.L. performed mass spectrometry. M.D. and D.W.D prepared frozen brain samples and performed immunohistochemistry staining. Q.C. and M.R.S. performed solvation energy calculation. All authors analyzed the results and wrote the manuscript. D.S.E. supervised the project.

## Data availability

Cryo-EM maps and atomic models of FTLD-TDP donor 1 have been deposited into the Worldwide Protein Data Bank (wwPDB) and the Electron Microscopy Data Band (EMDB) with accession codes PDB 7SAQ and EMD-24953 for PM1, PDB 7SQR and EMD-24954 for PM2, and PDB 7SAS and EMD-24955 for PM3. Mass spectrometry data has been deposited to the ProteomeXchange Consortium via MassIVE partner repository with the dataset identifier PXD029876. Any other relevant data are available from the corresponding author upon reasonable request.

## Methods

### Post-mortem human brain samples

The Brain Bank for Neurodegenerative Disorders at Mayo Clinic Florida provided post-mortem brain tissues from patients with neuropathologically confirmed FTLD-TDP. Information on human donors is provided in Table 1. Autopsies were performed after consent by the next-of-kin or someone with legal authority to grant permission. The brain bank operates under protocols approved by the Mayo Clinic Institutional Review Board.

### Immunohistochemistry staining

Post-mortem brains were immersion-fixed in 10% formalin, and sections of brain were embedded in paraffin, cut on a microtome at 5μm thickness and mounted on positively-charged glass slides. Sections were dried overnight and used for immunohistochemistry staining. Paraffin-embedded brain sections were deparaffinized in xylene, and rehydrated through a series of ethanol solutions, followed by washing in deionized H_2_O. Antigen retrieval was performed by steaming slides in deionized H_2_O or Tris-EDTA (DAKO), pH 9.0 for 30 minutes followed by a 5-minute incubation in DAKO Peroxidase Block (DAKO, Catalog No. S2001) to block endogenous peroxidase activity. Slides were blocked with DAKO Protein Block Serum-Free (DAKO, Catalog No. X0909) for 1 hour, and incubated with Anti TAR DNA-Binding Protein 43 (TDP-43) phospho-Ser409/410 mAb (Cosmo Bio USA, Catalog No. CAC-TIP-PTD-M01, Lot No. 11-9-20) diluted 1:1000 for 45 minutes. After washing, sections were incubated for 30 minutes in DAKO Envision-Plus System HRP Labelled Polymer Anti-Mouse (DAKO, Catalog No. K4001). Peroxidase labeling was visualized with the Liquid DAB+ Substrate Chromogen System (DAKO, Catalog No. K3468). Glass mounted tissue sections were viewed with an Olympus BX51 and microscopic images were captured with Olympus DP73 digital camera.

### Fibril extraction from FTLD-TDP patient brains

Frozen brain tissues were weighed, diced into small pieces, and resuspended in 10 mL/gram of homogenization-solubilization (HS) buffer (20 mM Tris-HCl, pH 7.5, 150 mM NaCl, 0.1 mM EDTA, 1 mM dithiothreitol) supplemented with 1:100 (v/v) Halt protease inhibitor (Thermo Scientific). Resuspended tissue was homogenized using a Polytron homogenizer (Thomas Scientific) for 45 seconds and mixed with 10 mL/gram of 4% (w/v) N-lauroyl-sarcosine (sarkosyl, Sigma), 2 U μl^-1^ homemade Benzonase and 4 mM MgCl_2_. Mixed solution was incubated at 37 °C with constant shaking at 300 r.p.m. for 45 minutes. Sample solution was then mixed with 10 mL/gram of ice-cold HS buffer with 0.5% (w/v) sarkosyl, and centrifuged at 3,000 × g at 4 °C for 5 minutes. The supernatant was extracted and centrifuged at 21,000 × g for 30 minutes. The supernatant was discarded, and the pellet was resuspended in 1.5 mL/gram of Tris buffer (20 mM Tris-HCl, pH 7.5, 150 mM NaCl) and incubated at 4 °C overnight. After incubation, the solution was centrifuged at 6,000 × g for 5 minutes. The supernatant was transferred into a new tube, mixed with 1 % (w/v) sodium dodecyl sulfate (Invitrogen), and gently rotated at room temperature for 15 minutes. The solution was then sonicated for 3 minutes and centrifuged at 6,000 × g for 10 minutes. The supernatant was centrifuged at 21,000 × g for 30 minutes, and the pellet was resuspended in 10 μL/gram of Tris buffer, sonicated for 3 minutes and used for EM. Approximately 1 gram of brain tissue from each patient was used for cryo-EM study.

Fibrils were also extracted using ice-cold water following a previously described protocol^33^. Briefly, brain tissues were diced and resuspended in 4 mL/gram of Tris-calcium buffer (20 mM Tris, pH 8.0, 138 mM NaCl, 2 mM CaCl_2_, 0.1% NaN_3_), then centrifuged at 3100 × g at 4 °C for 5 minutes. The supernatant was collected and the Tris-calcium buffer wash was repeated four more times. After the fifth wash, the pellet was resuspended in Tris-calcium buffer with collagenase and incubated at 37°C with constant shaking at 500 r.p.m. overnight, then centrifuged at 3100 × g at 4 °C for 30 minutes. The pellet was resuspended in 4 mL/gram of Tris-EDTA buffer (20 mM Tris, pH 8.0, 140 mM NaCl, 10 mM EDTA, 0.1% NaN_3_), then centrifuged at at 3100 × g at 4 °C for 5 minutes. The supernatant was collected and the Tris-EDTA buffer wash was repeated nine more times. After the tenth wash, the pellet was resuspended in 2 mL/gram of ice-cold water, then centrifuged at 3100 × g at 4 °C for 5 minutes. The fibril-containing supernatant was collected and the ice-cold water extraction was repeated nine more times. Fibrils extracted using sarkosyl and ice-cold water exhibited similar morphologies when examined by negative stain EM (Fig. 1d), suggesting that sarkosyl does not induce the fibril formation during the extraction process. Fibril samples extracted using sarkosyl were more abundant and contained less contaminants, thus were used for cryo-EM study.

### Negative stain transmission electron microscopy (TEM)

400-mesh carbon-coated formvar support films mounted on copper grids (Ted Pella, Inc.) were glow-discharged for 30 seconds before sample preparation. 3 μL of sample solution was applied to the grids and incubated for 3 minutes, then excess sample solution was blotted off using filter paper. Grids were stained with 3 μl of 2% uranyl acetate (Electron Microscopy Sciences) for 1 minute, washed with an additional 3 μl of 2% uranyl acetate and air-dried for 2 minutes. The grids were imaged using a Tecnai T12 transmission electron microscope (FEI).

### Cryo-EM data collection and processing

To prepare the cryo-EM grids, we applied 2.6 μl of sample solution onto Quantifoil 1.2/1.3 200 mesh electron microscope grids glow-discharged for 2 minutes before use. Grids were plunge frozen into liquid ethane using a Vitrobot Mark IV (FEI). Cryo-EM data were collected on a Titan Krios transmission electron microscope (FEI) equipped with a K3 Direct Detection Camera (Gatan), operated with 300 kV acceleration voltage and energy filter width of 20 eV. Super-resolution movies were collected with a nominal physical pixel size of 1.1 Å/pixel (0.55 Å/pixel in super-resolution movie frames) and a dose per frame of ~1 e-/Å^2^. A total of 36 frames with a frame rate of 12 Hz were taken for each movie, resulting in a final dose of ~36 e-/Å^2^ per image. Manual data collection was performed using Leginon software package^34^. Cryo-EM datasets of four FTLD-TDP donors were collected and processed separately.

For the dataset of donor 1, particle picking was performed using CrYOLO^35^, trained by ~150 micrographs that we picked manually. Particle extraction, two-dimensional classification, helical reconstruction, and 3D refinement were performed in RELION^36,37^. Particles were extracted using an inter-box distance of 102.4 Å and a box size of 1024 pixels scaled down to a 432 pixels box size. 2D classification with tau_fudge 2 was performed with all particles, and we found that all identifiable classes could be grouped into one of three fibril polymorphs, which we named PM1 (78.1% of all particles), PM2 (17.8% of all particles) and PM3 (4.1% of all particles). No other fibril morphologies were identified in this dataset. Particles from each polymorph were selected and used for 3D classification with K=3 (for PM1) or K=1 (for PM1 and PM2), using a Gaussian cylinder as the initial model. The best 3D classes were used as the initial model for subsequent 3D classifications with smaller box size particles. We re-extracted particles from all micrographs using an inter-box distance of 32 Å and a box size of 686 pixels scaled down to a 320 pixels box size. We manually selected 686-pixel particles for each polymorph based on the 2D classification of 1024-pixel particles. For example, if a 1024-pixel particle was classified as PM1 in 2D classification, then all 686-pixel particles extracted from that particular filament were selected for the PM1 polymorph. These manually selected particles were used for 2D classification for each polymorph separately, and the best 2D classes were selected for 3D classification. Two rounds (for PM1) or one round (for PM2 and PM3) of K=3 3D classification was performed. Particles from the best 3D class for each polymorph were re-extracted using an inter-box distance of 32 Å and a box size of 320 pixels (no scaling) and used for high-resolution 3D refinement. To improve resolution, additional 2D classifications were performed for PM1 and PM2, with tau_fudge starting at 2 and increased incrementally to 8 in the final iterations. The 2D classes with clear 4.8 Å separation were selected for further 3D refinement. High-resolution gold-standard refinement was performed for each polymorph and the initial near-atomic resolution maps with refined helical parameters were generated. CTF refinement and Bayesian polishing was performed, and the final reconstructions were generated by one or two rounds of additional golden-standard refinement. The resolution of each reconstruction was estimated using the 0.143 Fourier shell correlation (FSC) resolution cutoff. See Table 2 for data collection and processing statistics for donor 1.

For the datasets of donor 2-4, similar strategies were applied for data collection and processing. All three datasets were processed independently up to and throughout 2D classification, with no information from other dataset introduced. 2D classification of all particles suggested that these three datasets also contain PM1, PM2, and PM3, with similar morphology and distribution as donor 1 (Fig. S4), and no other identifiable fibril morphologies. During initial 3D reconstruction, we processed the donor 2 dataset independently, whereas for donor 3 and 4 we used the maps generated from the donor 1 dataset as initial model to expedite data processing. We believe these reconstructions were not biased because we used a 30 Å low-pass-filter on the initial model so that any higher-resolution information beyond 30 Å would originate from the dataset and not the initial model, and the final map of each polymorph was near-atomic (3.5-5.3 Å). After initial 3D reconstruction, a similar strategy was applied for each dataset to generate the final maps. See Table S2 for data collection and processing statistics for donors 2-4.

### Atomic model building

Our first model building efforts focused on PM1 of donor 1. We attempted to model the sequence of TDP-43 in the map using two methods: manual building with Coot^38^ and automatic building using phenix.sequence_from_map^39^. No satisfactory model could be made with the TDP-43 sequence; multiple sidechains were inconsistent with densities even taking into consideration the potential PTMs such as phosphorylation of serine residues (i.e. the best model shown in Fig. S7a).

We then applied an unbiased strategy for model building: we built two poly-alanine backbones with opposite directionality of N- and C-termini. For each model, we mutated the residues to amino acids that best fit the side chain densities (Fig. S7a). The sequences of two resulting models were used as queries in BLAST to identify the protein in our maps. We restricted the BLAST database to human sequences (taxid 9606), knowing that the brain samples that produced the fibrils were human. No other parameters of the search were altered from the default values. TMEM106B stood out as the only significant match to our query sequence.

We obtained five hits in total, but all five hits were sequences of the same protein, TMEM106B (including the protein labeled as hypothetical protein FLJ11273) (Table S4). The percent identity of the hits was low (~16%), but the coverage was high, covering 125 of the 134 residues in the query (93%) without gaps or insertions in the alignment. All of these TMEM106B hits score above the default expect threshold value of 10. The “expect” value that we obtained for the highest scoring hit, TMEM106B, was 0.003. This value means the number of times that a match as good or better would occur by chance is 0.003 in a database of this size. We interpret this value to indicate that the match is significant. If we increase the permissiveness of the expect threshold to higher values, we find the next best hit (6^th^) yields an expect value of 224, indicating this sequence (nuclear receptor subfamily 2 group E member) is an insignificant match to our query.

The TMEM106B sequence was threaded onto the initial model with phenix.sequence_from_map. The TMEM106B model fit the density better than the query model (Fig. S7b). We extended the model to include 5 layers by applying the helical symmetry operators of the map and the model was refined with phenix.real_space_refine^40^. The orientations of mainchain oxygen atoms were manually adjusted to form inter-layer hydrogen bonds. The model was refined again with phenix.real_space_refine and the final model was validated using MolProbity^41^. The model was built and adjusted with Coot^38^.

For PM2 and PM3 of donor 1, we fit a single layer of PM1 model into PM2 or PM3 maps with rigid body fit, and manually adjusted the model with Coot. A five-layer model (containing 10 chains) was generated with the helical symmetry of PM2 or PM3. Model refinement was performed with phenix.real_space_refine and the final model was validated using MolProbity. No major conformational changes were observed during model refinement.

For all polymorphs of donor 2-4, we did not build models *de novo*. Instead, we fit a single chain of each donor 1 polymorph into corresponding donor 2-4 maps as a rigid body (Fig. S6). For PM2 and PM3, a one-layer model was generated by applying their helical symmetry. All models fit the maps well, suggesting that TMEM106B forms fibrils with the same fold in different FTLD-TDP patients.

### Standard free energy calculation

Standard free energy values were estimated using a solvation energy algorithm as described previously^42^. We note that our free energy calculations neglect the contribution of glycans, as we cannot confirm the composition or conformation of the sugar groups.

### Patient genotyping

Genomic DNA was extracted from patient brain tissues by the UCLA Technology Center for Genomics & Bioinformatics. Primers were designed to amplify the 7 exonic DNA fragments encoding TMEM106B (hg38 chr7:12,214,811-12,231,975, exons 3 to 9); primers were synthesized by Integrated DNA Technologies. PCR amplification was performed using Phusion High-Fidelity DNA Polymerase (New England Biolabs) and the PCR products were sequenced by Genewiz. For donor 3, sequencing results suggest the genotype of TMEM106B is wild type. For donors 1, 2 and 4, the sequencing results indicate a heterozygous Thr185Ser variant: at position hg38 chr7:12,229,791 in exon 6 of the TMEM106B gene, there is equal detection of the wildtype cytosine nucleotide and the variant guanine nucleotide. This single nucleotide polymorphism would induce a point mutation from wildtype Thr185 (ACC) to variant Ser185 (AGC). From the 50-50 distribution of cytosine and guanine nucleotides measured in the sequencing chromatogram, we conclude that one allele contains the wildtype gene and the other harbors the variant.

### Western blotting

The supernatant and pellet from the first 21,000 × g centrifugation step of fibril extraction were used as samples for immunoblotting. Sample solutions were mixed with SDS-PAGE loading dye containing 2M urea and 1M β-mercaptoethanol, sonicated for 10 minutes in ice water, then boiled at 100 °C for 10 minutes. Samples were loaded onto a NuPAGE 4-12%, Bis-Tris, 1.0 mm, 12-well Mini Protein Gel (Invitrogen) and electrophoresis was performed using 200 V for 30 minutes. Proteins were wet transferred onto 0.2 μm nitrocellulose membranes (Bio-Rad) by application of a constant 35 V overnight, in transfer buffer consisting of 25mM Tris, pH8.3, 192mM glycine, 20% (w/v) methanol. Membranes were incubated with gentle rocking in 5% (w/v) Blocking-Grade Blocker (milk, Bio-Rad) in 1X Tris-buffered saline, with 0.1% (v/v) Tween20 (TBST). Membranes were incubated for 1 hour with TMEM106B Antibody (Novus Biologicals, Catalog No. NBP1-91311, Lot No. QC18333-42825) diluted 1:300, TDP-43 Monoclonal Antibody (Proteintech, Catalog No. 60019-2-IG, Lot No. 10011784) diluted 1:500, or Anti TAR DNA-Binding Protein 43 phospho-Ser409/410 mAb (Cosmo Bio USA, Catalog No. CAC-TIP-PTD-M01, Lot No. 11-9-20) diluted 1:300 in 2% milk in TBST. Membranes were washed three times in TBST with gentle rocking for 5 minutes. Then, membranes were incubated for 1 hour with either Goat Anti-Rabbit IgG HRP (Invitrogen, Catalog No. A27036, Lot No. 2116291) or Goat Anti-Mouse IgG HRP (Abcam, Catalog No. ab205719, Lot No. GR3271082-2) diluted 1:4000 in 2% milk in TBST. Membranes were washed three times in TBST with gentle rocking for 5 minutes. Pierce ECL Plus Western Blotting Substrate (Thermo Scientific) was applied to membranes. Membranes were imaged using an Azure 600 (Azure Biosystems).

In addition to the ~35kDa band on the TMEM106B western blot (Fig. 1b, left), bands with molecular weights (>110kDa) higher than what is expected for TMEM106B were observed in the sarkosyl-soluble (three bands) and insoluble fractions (one band). These bands could correspond to different cleavage products or different post translational modification states of TMEM106B. The high molecular weights could be due to TMEM106B oligomerization or association with other macromolecules to form SDS-resistant complexes. Although the identify of these species cannot be confirmed, the conclusion of this study is unaffected as the ~35kDa band has been validated by mass spectrometry.

### Immunogold labeling

400-mesh carbon coated copper grids were glow discharged for 30 seconds before sample preparation. 3 μL of sample solution was applied to the grids and incubated for 3 minutes, then excess sample solution was blotted off using filter paper. Blocking buffer (phosphate buffered saline, pH 7.4, 0.1% w/v gelatin) was applied to the grids and incubated for 10 minutes; excess solution was blotted off. TMEM106B antibody specific for residues 2-53 (Atlas Antibodies, Catalog No. HPA058342, Lot No. 22721), residues 204-253 (Novus Biologicals, Catalog No. NBP1-91311, Lot No. QC18333-42825) or residues 218-252 (antibodies-online Inc., Catalog No. ABIN6578799, Lot No. SA160811DF), or TDP-43 Monoclonal Antibody (Proteintech, Catalog No. 60019-2-IG, Lot No. 10011784) diluted 1:100 in blocking buffer was applied to the grids and incubated for 30 minutes; excess solution was blotted off. Grids were washed five times with blocking buffer, and excess liquid was blotted off between each wash. 6 nm Colloidal Gold AffiniPure Goat Anti-Mouse IgG (Jackson ImmunoResearch Laboratories, Inc., Catalog No. 115-195-146, Lot No. 150545) or 6 nm Colloidal Gold AffiniPure Goat Anti-Rabbit IgG (Jackson ImmunoResearch Laboratories, Inc., Catalog No. 111-195-144, Lot No. 146470) diluted 1:8 in blocking buffer was applied to the grids and incubated for 30 minutes; excess solution was blotted off. Grids were washed five times with distilled water, and excess liquid was blotted off between each wash. Finally, grids were stained with 3 μl of 2% uranyl acetate for 1 minute, washed with an additional 3 μl of 2% uranyl acetate and air-dried for 10 minutes. The grids were imaged using a Tecnai T12 electron microscope.

### Mass spectrometry

The pellet from the first 21,000 × g centrifugation step of fibril extraction was subjected to SDS-PAGE as previous described. Gels were stained by InstantBlue Coomassie Protein Stain (Abcam) and gel bands were excised for mass spectrometry analysis. Protein digestion and peptide identification were adapted from a previously described protocol^43,44^. Briefly, proteins entrapped in gel bands were reduced with 10 mM dithiothreitol (Sigma) at 60°C for 1 hour, alkylated with 50 mM iodoacetamide (Sigma) at 45°C for 45min in the dark, digested with 200ng trypsin (Promega) at 37°C overnight and then 100ng GluC (New England BioLabs) was added for another overnight incubation at 25°C. Peptides were extracted from the gel bands in 50% acetonitrile/49.9% water/ 0.1% trifluoroacetic acid (TFA) and cleaned with C18 StageTip^45^ before mass spectrometry analysis. Digested peptides were separated on an EASY-Spray column (25cm x 75μm ID, PepMap RSLC C18, 2μm, Thermo) connected to an U3000 RSLCnano HPLC (Thermo) and eluted using a gradient of 3%-32% acetonitrile in 0.1% formic acid and a flow rate of 300nL/min (total time 45 minutes). Tandem mass spectra were collected in a data-dependent manner with an Orbitrap Exploris 480 mass spectrometer (Thermo) interfaced to a nano-ESI source (Thermo). Raw MS/MS data were analyzed using Mascot (version 2.5) and files were searched against the UniProt human database (as of December 19, 2018; 20,387 entries) supplemented with common laboratory contaminants. The search parameters were: enzyme specificity: trypsin and GluC; maximum number of missed cleavages: 6, precursor mass tolerance, 10ppm; product mass tolerance, 0.02 Da; variable modifications included cysteine carbamidomethylation and methionine oxidation. PSMs were filtered to 1% false discovery rate using the target-decoy strategy. Additional searches were performed applying identical parameters to Mascot’s error-tolerant algorithm. The mass spectrometry proteomics data has been deposited to the ProteomeXchange Consortium via MassIVE partner repository with the dataset identifier PXD029876.

Mass spectrometric analysis identified peptide LNNISIIGPLDMK (corresponding to TMEM106B 181-193) from the sarkosyl-insoluble fraction of donor 1. The Thr185Ser variant is present in this peptide, suggesting the variant protein is present in the fibrils. The peptide also contains the N184 glycosylation site without a glycan modification. Detection of this unglycosylated peptide does not contradict the glycosylation of N184 suggested by the cryo-EM structures. The type and size of the glycan could not be determined, thus it was much more difficult to identify the glycosylated peptide using mass spectrometry and only the unglycosylated peptide was detected.

---

**Fig. S1.**
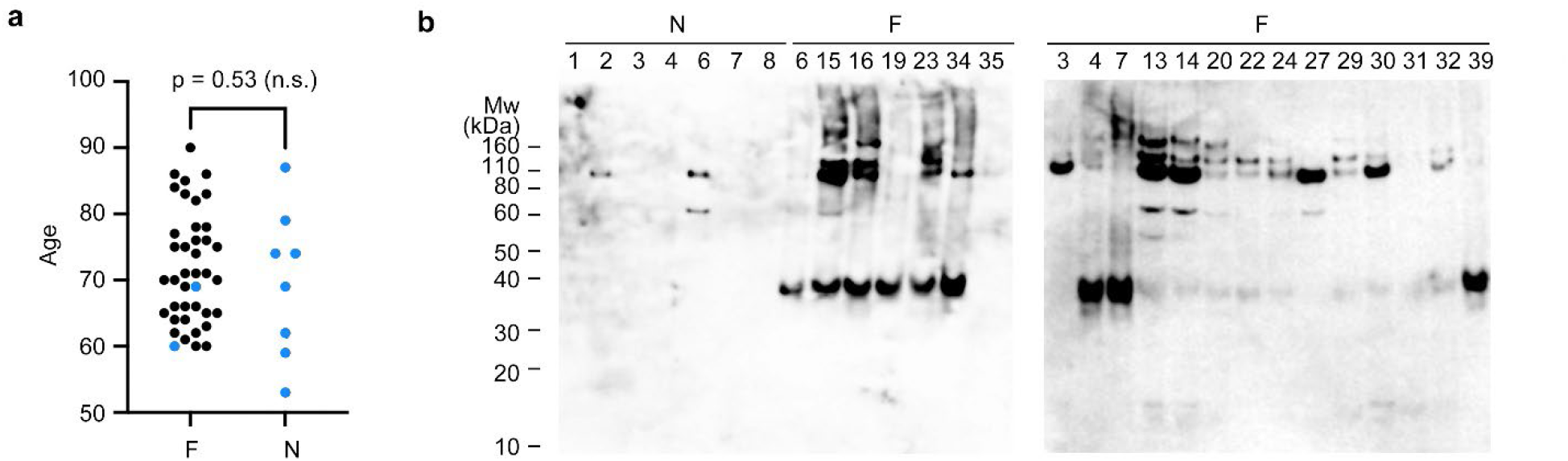
Comparison of FTLD-TDP and non-FTLD-TDP donors. **a,** Age distribution of FTLD-TDP donors (F) and non-FTLD-TDP controls (N). Donors with and without fibrils detected under negative stain EM are colored black and blue, respectively. P-value of 0.53 (n.s., not significant) from an unpaired t-test suggests that the presence of fibrils is disease-dependent but not age-dependent. **b,** Western blots of sarkosyl-insoluble fractions from FTLD-TDP (F) and non-FTLD-TDP donors (N) probed by TMEM106B antibody. The ~35 kDa TMEM106B-positive band was found in none of the non-FTLD-TDP donors (N5 shown in Fig. 1b as donor 5) and all of the FTLD-TDP donors except F35, F3, and F27. Fibrils were not observed in F3 and F27 by negative stain EM, consistent with the western blot. Fibrils were observed in F35, which suggests that western blot may not always be accurate in detecting TMEM106B aggregation. Western blot membranes were exposed with equal time.

**Fig. S2.**
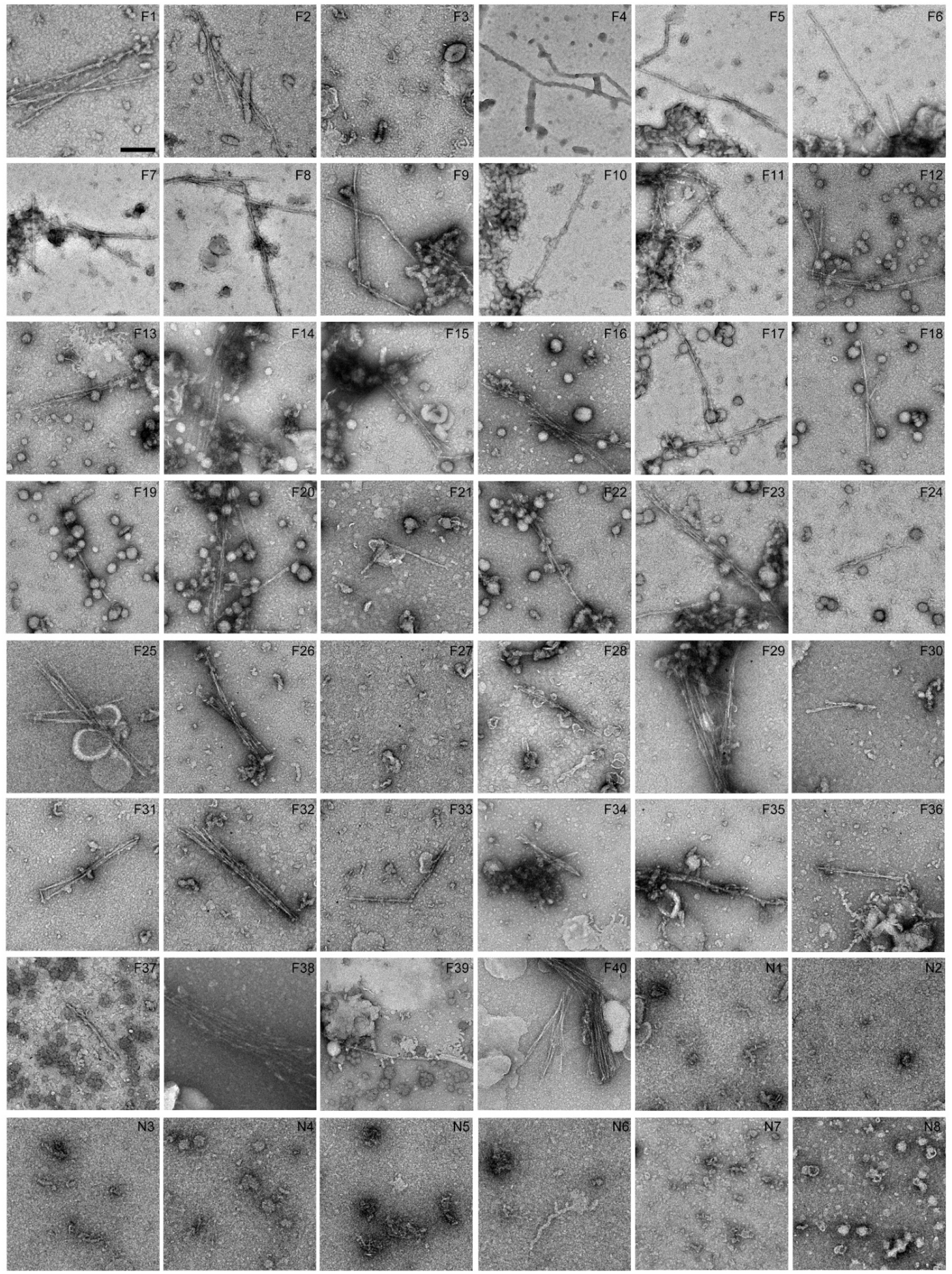
Fibril screen of FTLD-TDP and non-FTLD-TDP donors by negative stain EM. Negative stain EM images of sarkosyl-insoluble fractions from all donors. Donors 1-5 (F26, F36, F17, F40 and N5, respectively) are also shown in Fig. 1a. Fibrils with similar morphologies were found in all FTLD-TDP donors except F3 and F27. No fibrils were found in any non-FTLD-TDP donors.

**Fig. S3.**
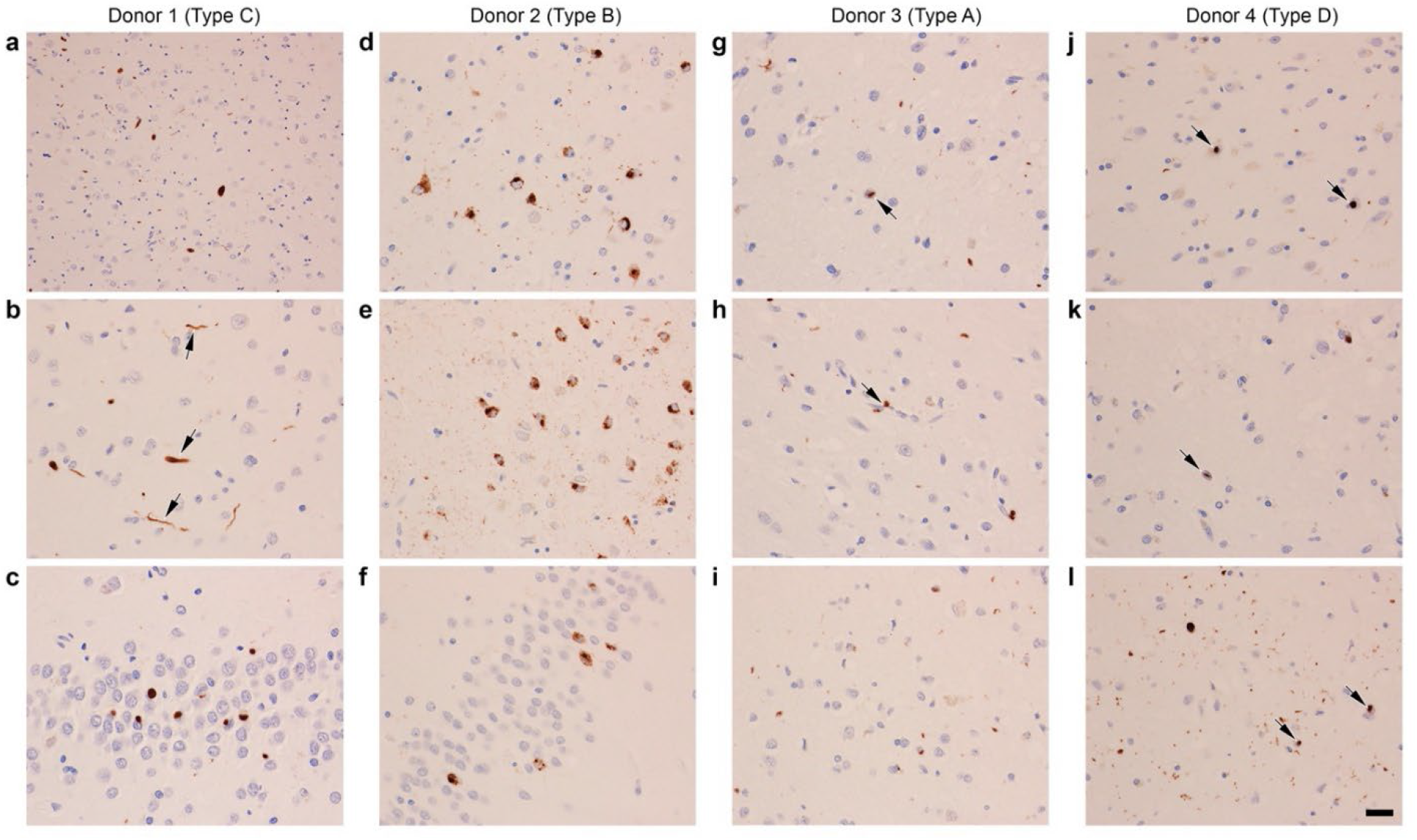
Neuropathological diagnosis of donors 1-4 as FTLD-TDP. Immunohistochemistry staining using a phosphor-Ser409/410 TDP-43 antibody was performed for brain sections from donors 1-4. All four donors were confirmed to be FTLD-TDP cases, representing the 4 subtypes A to D, respectively. For all figures, scale bar 20 μm. **a**, **b**, **c**, Donor 1 is FTLD-TDP type C, with long, thick neurites (arrows in **b**) and “Pick-body like NCI” in dentate fascia (**c**). **d**, **e**, **f**, Donor 2 is FTLD-TDP type B, displaying characteristic granular cytoplasmic NCI in cortex (**d**), hippocampus (**e**) and dentate fascia (**f**). **g**, **h**, **i**, Donor 3 is FTLD-TDP type A, exhibiting small dense neuronal cytoplasmic inclusions (NCI), sparse neuronal intranuclear inclusions (NII, arrow in **g**), and perivascular glial inclusions (arrow in **h**). **j**, **k**, **l**, Donor 4 is FTLD-TDP type D, shown by frequent NII (arrows in **j**, **k** and **l**) and small NCI and neurites. **a**, **b**, **d**, **e**, **g**, **h**, **i**, **j**, **k**, **l** are brain sections from the temporal cortex. **c** and **f** are brain sections from dentate fascia.

**Fig. S4.**
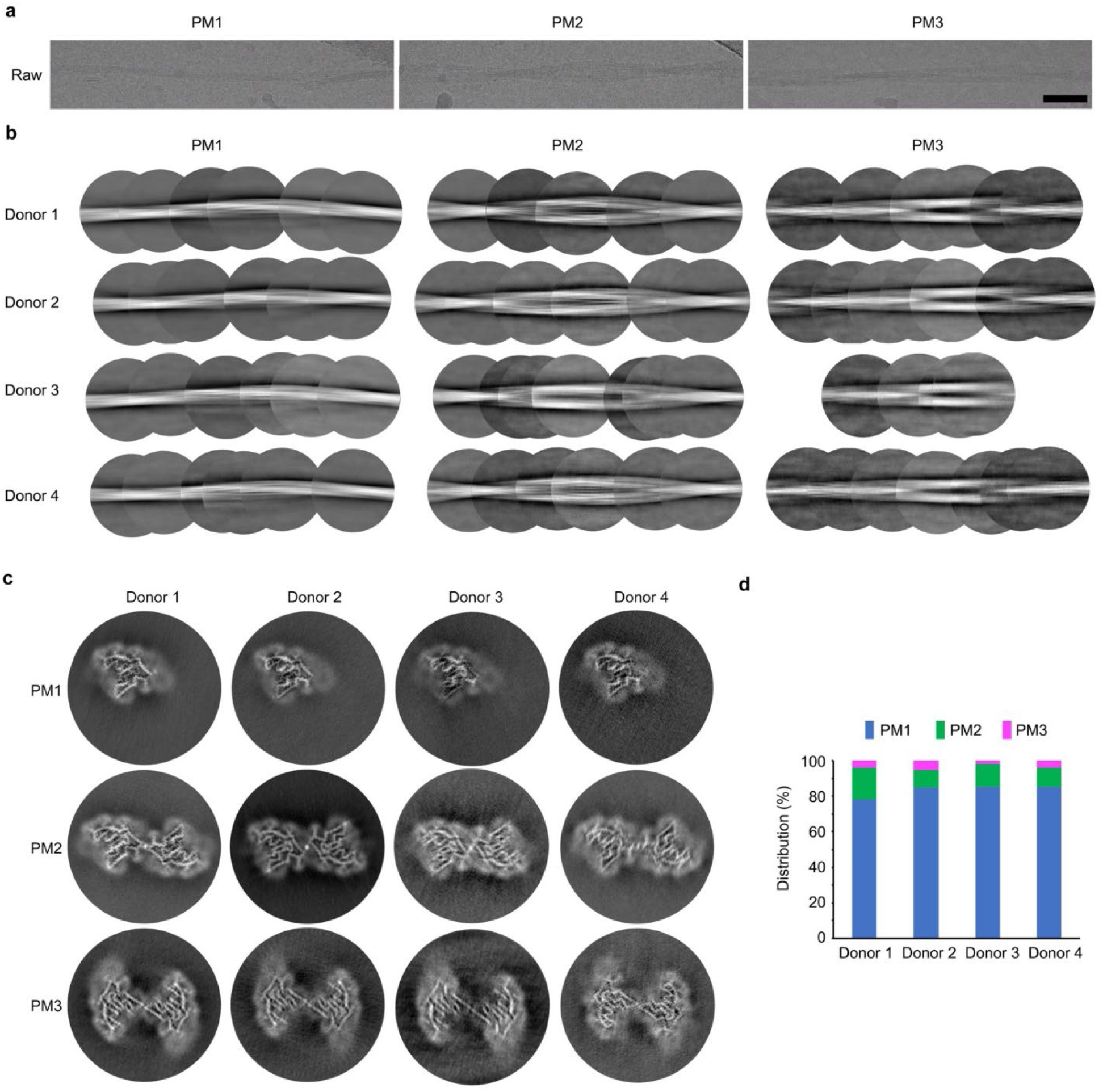
Cryo-EM data processing of amyloid fibrils from FTLD-TDP donors 1 to 4. **a**, Representative micrographs of PM1-3 from FTLD-TDP donor 1. Scale bar 500 Å. **b**, Representative 2D classes of PM1-3 from FTLD-TDP donors 1 to 4. 2D classes are stitched together to show a full cross-over of each morphology. **c**, 3D reconstructions of PM1-3 from the four FTLD-TDP donors. **d**, Distributions of the three fibril polymorphs in the four FTLD-TDP donors.

**Fig. S5.**
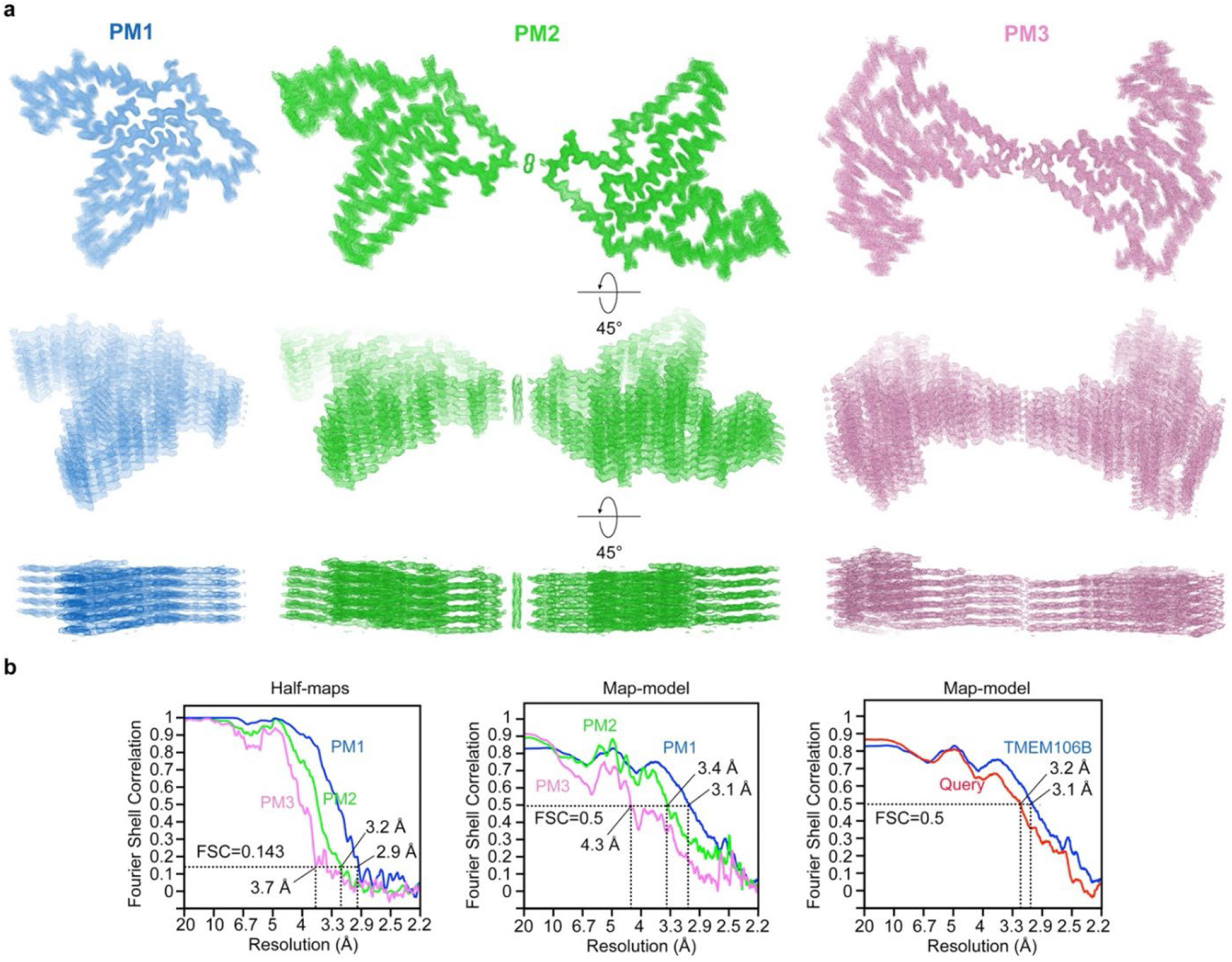
Cryo-EM maps and FSC curves of FTLD-TDP donor 1. **a**, Different views of the cryo-EM maps from FTLD-TDP donor 1 with five layers shown. **b**, FSC curves between two half-maps (left) and the cryo-EM reconstruction and refined atomic model (middle) of each polymorph PM1 (blue), PM2 (green), and PM3 (pink). FSC curves between cryo-EM reconstruction and the query model (red, Direction 1 chain of Fig. S5a) and the atomic model of PM1 from FTLD-TDP donor 1 (blue) are compared on the right.

**Fig. S6.**
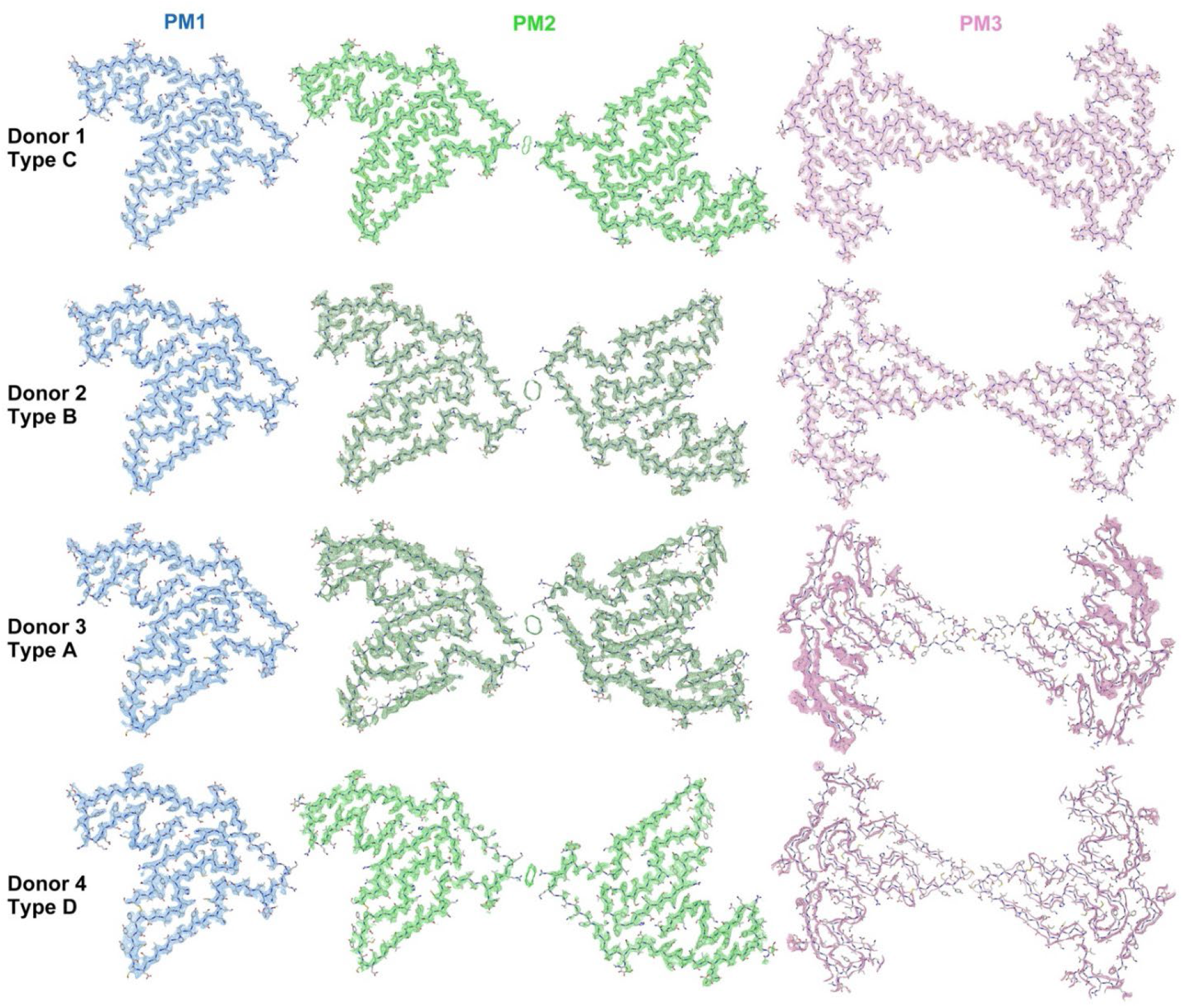
Cryo-EM maps of TMEM106B fibrils from FTLD-TDP donors 1 to 4. The models of PM1-3 from donor 1 were rigid body fitted into the maps of PM1-3 from donors 2 to 4. These cryo-EM structures reveal the polymorphs from all four donors share the same protofilament fold. Two subtypes of PM2 are exhibited by donors 1 and 4 (light green) and donors 2 and 3 (dark green), respectively.

**Fig. S7.**
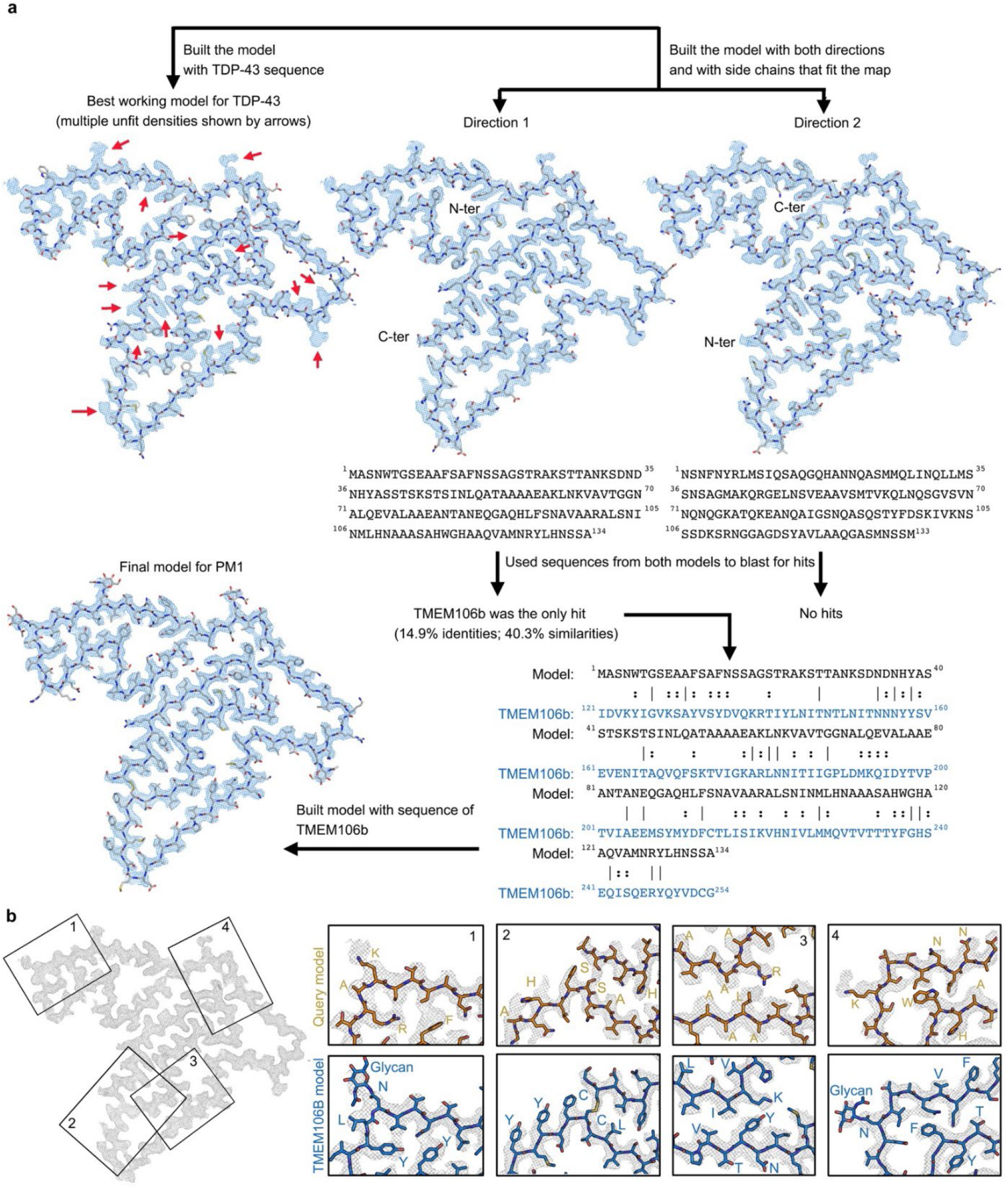
Atomic model building. **a,** Model building flowchart for PM1 of FTLD-TDP donor 1. In the sequence alignment (bottom right), lines indicate identical residues and two dots indicate similar residues. **b,** Comparison of query model (Direction 1 chain, orange) with final model built with TMEM106B sequence (blue). Four representative regions of the cryo-EM map are shown. Residues that show clear differences in side chain density fitting are labeled in both models.

**Fig. S8.**
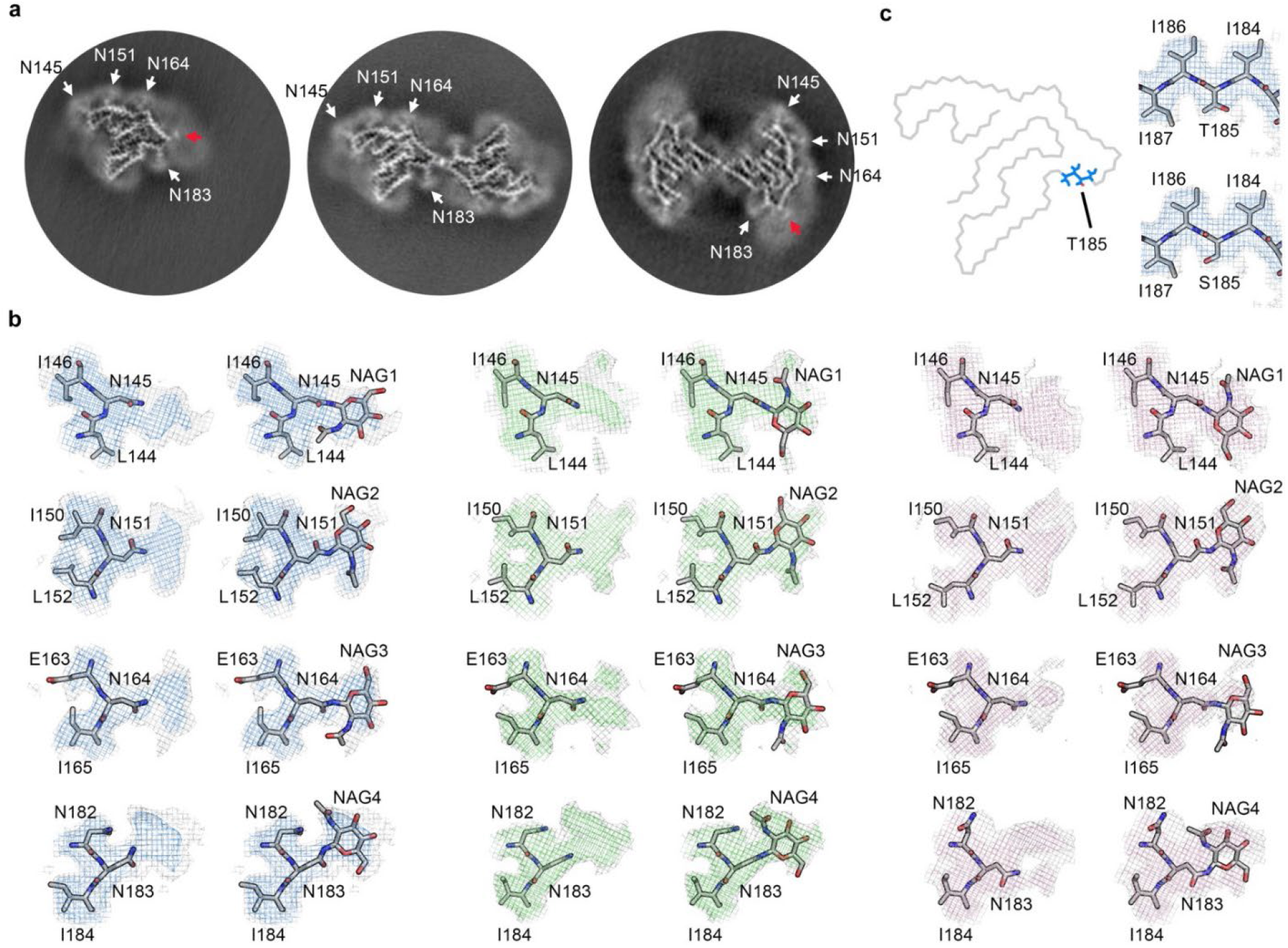
Structural analysis of TMEM106B fibrils. **a**, Three-dimensional reconstructions of PM1-3 from FTLD-TDP donor 1. White arrows point to the four glycosylation sites within the fibril core. Red arrows point to the additional densities outside the fibril core in PM1 and PM3, which may correspond to the binding of the same undefined, negatively-charged ligand present in the PM2 dimer interface (Fig. S8). **b**, Maps and models of the four glycosylation sites with or without the sugar group in PM1 (left), PM2 (middle), and PM3 (right) from FTLD-TDP donor 1. **c**, Position of Thr/Ser185 in the conserved fibril fold (left) and the map and model of the Thr/Ser185 environment (right, represented by PM1 from donor 1).

**Fig. S9.**
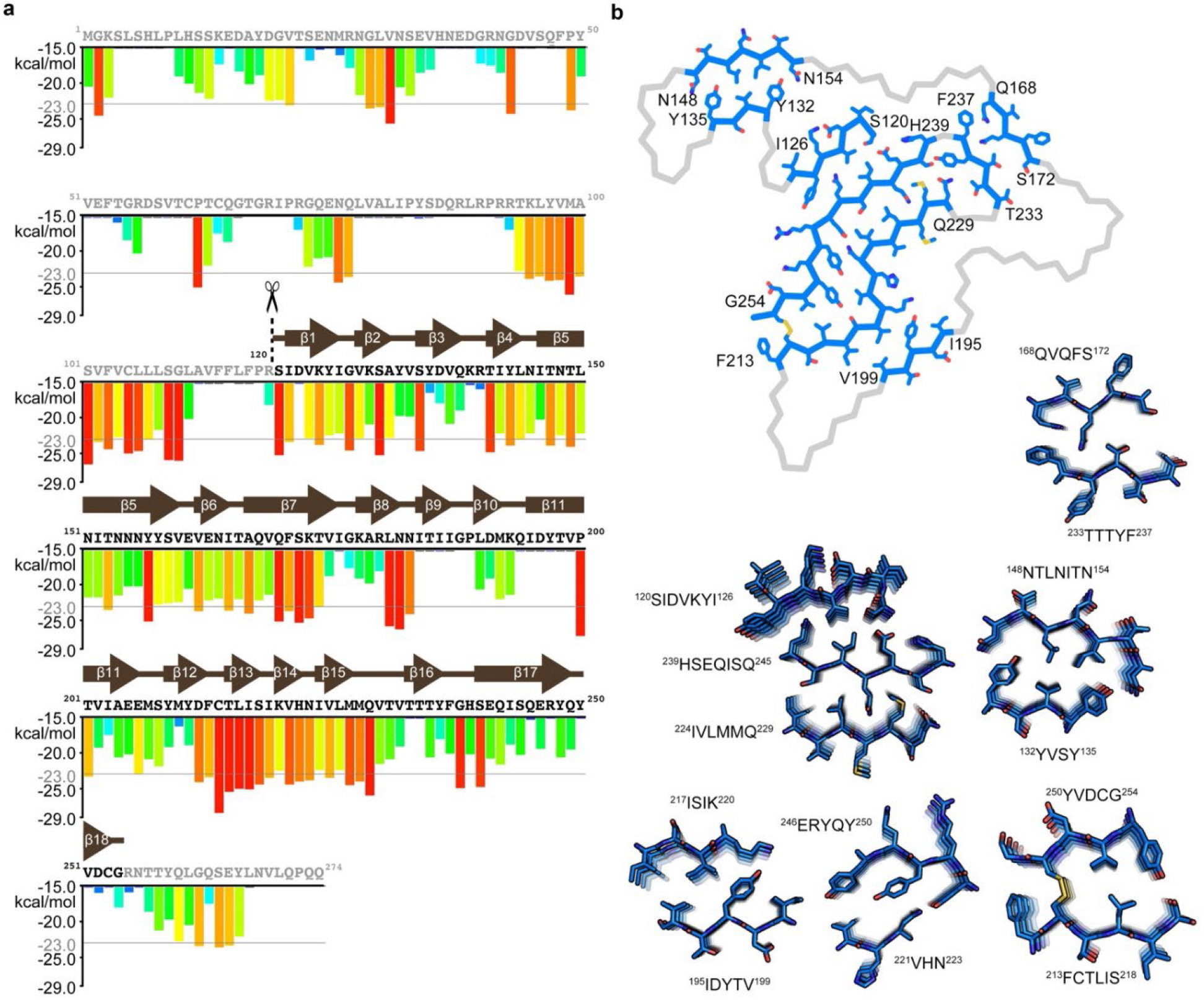
Steric zippers of TMEM106B fibrils. **a**, Amino acid sequence of TMEM106B and ZipperDB prediction. Red bars indicate high propensity for the six-residue fragments to form homotypic steric zippers. Residues that form β-strands in the TMEM106B protofilament core are designated by arrows. **b**, Heterotypic steric zippers in the conserved protofilament core of TMEM106B fibrils, represented by PM1 from donor 1.

**Fig. S10.**
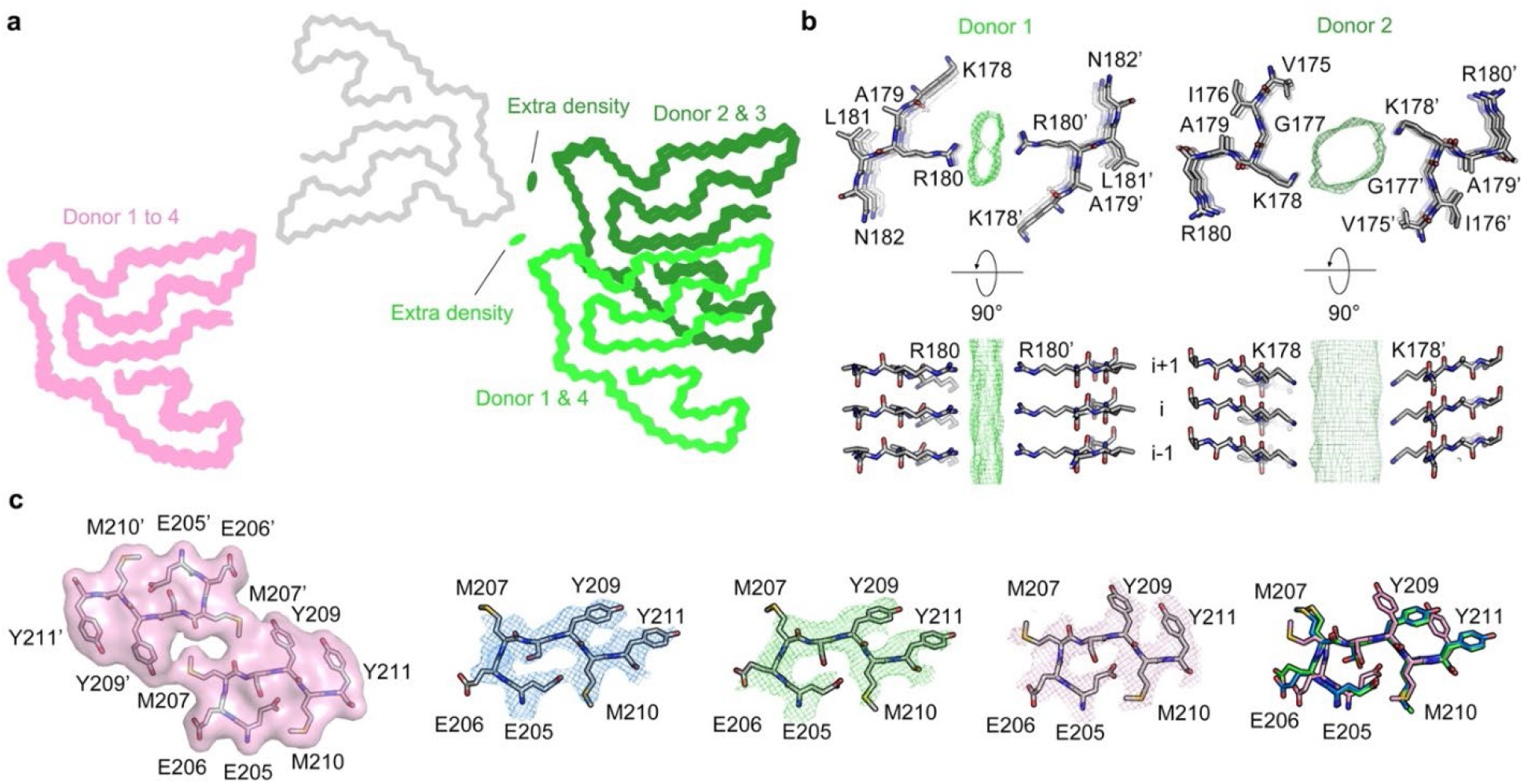
Dimer interfaces of PM2 and PM3 of TMEM106B fibrils. **a,** Dimer arrangements of PM2 and PM3 from FTLD-TDP donors 1 to 4. PM2 and PM3 from all donors are aligned at chain A (grey, represented by PM2 of donor 1), and chain B of PM2 (light green for donors 1 and 4, dark green for donors 2 and 3) and PM3 (pink) from each donor is shown. Residual densities in the PM2 dimer interfaces are shown as green ovals. The dimer arrangement of PM3 is consistent among all donors, whereas there are two subtypes of dimer arrangements for PM2. **b**, Atomic model and the additional density in the dimer interface of donor 1 (left, represents donors 1 and 4) or donor 2 (right, represents donors 2 and 3). In donor 1 and 4, Arg180 from each protofilament is on the opposite sides of an extra density in the middle of the PM2 dimer interface; in donor 2 and 3, the dimer interface is shifted to Lys178. Although two slightly different dimer interfaces were observed, we consider PM2 in all four FTLD-TDP donors to be the same morphology because of the similarity in dimer formation (see Discussion). **c**, Comparison of the residues near the PM3 interface (far left) from PM1 (blue), PM2 (green), PM3 (pink), and the superimposition of those residues from PM1-3 (far right). PM1-3 are all represented by FTLD-TDP donor 1.

**Fig. S11.**
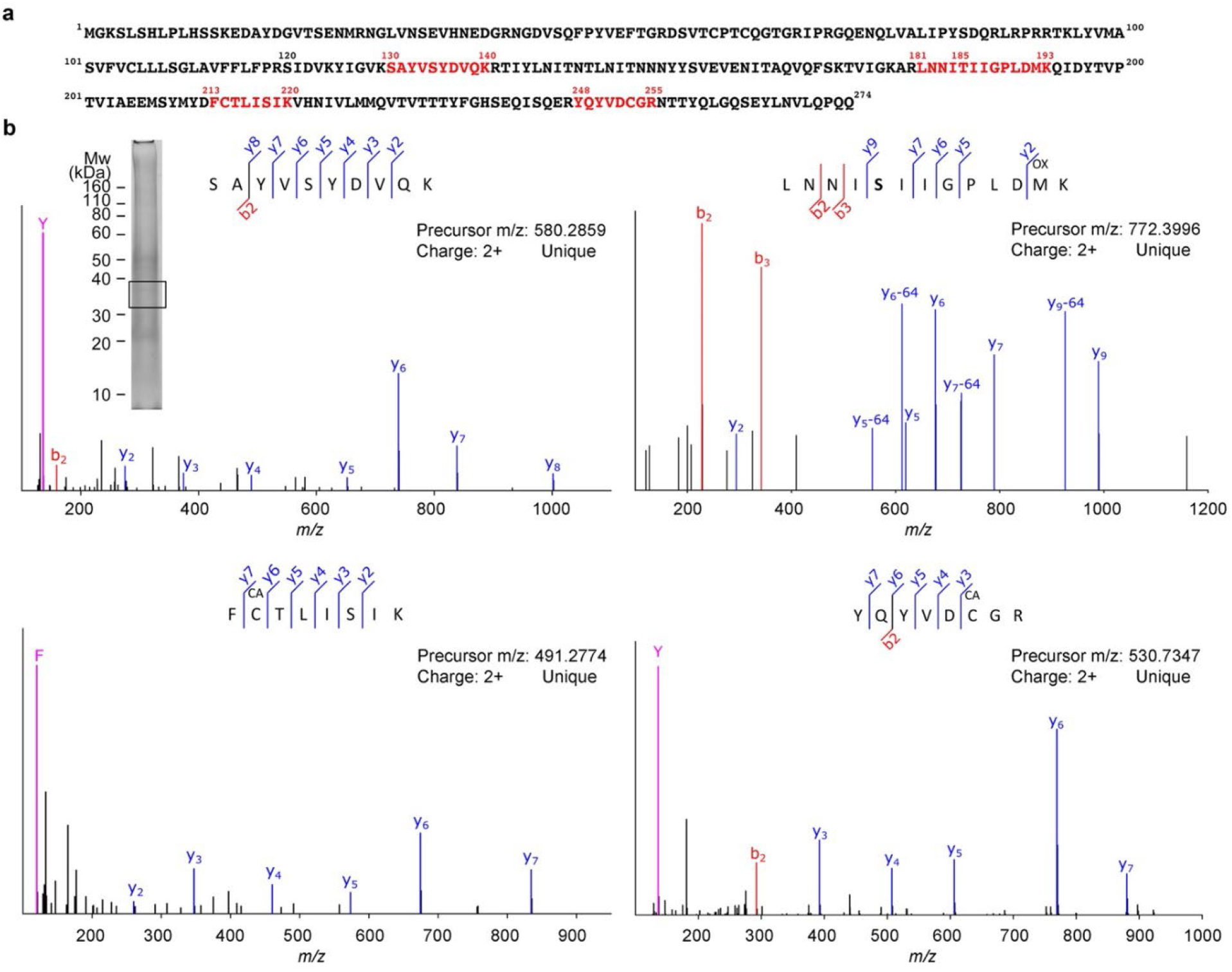
Detection of TMEM106B peptides by mass spectrometry. **a,** Sequence map of TMEM106B. Sequences in red indicate unique peptides detected by LC-MS/MS from excised gel bands. **b,** Fragmentation spectra (MS/MS) of detected peptides from TMEM106B. Detected fragment ions (b, y, and immonium ions) are labeled accordingly. The peptide modifications methionine oxidation (OX) and cysteine carbamidomethylation (CA) were observed. SDS-PAGE gel of sarkosyl-insoluble fraction of donor 1 shown as an insert. Box indicates the gel region, which corresponds to the ~35kDa band from the TMEM106B western blot (Fig. 1b), excised for LC-MS/MS analyses.

**Fig. S12.**
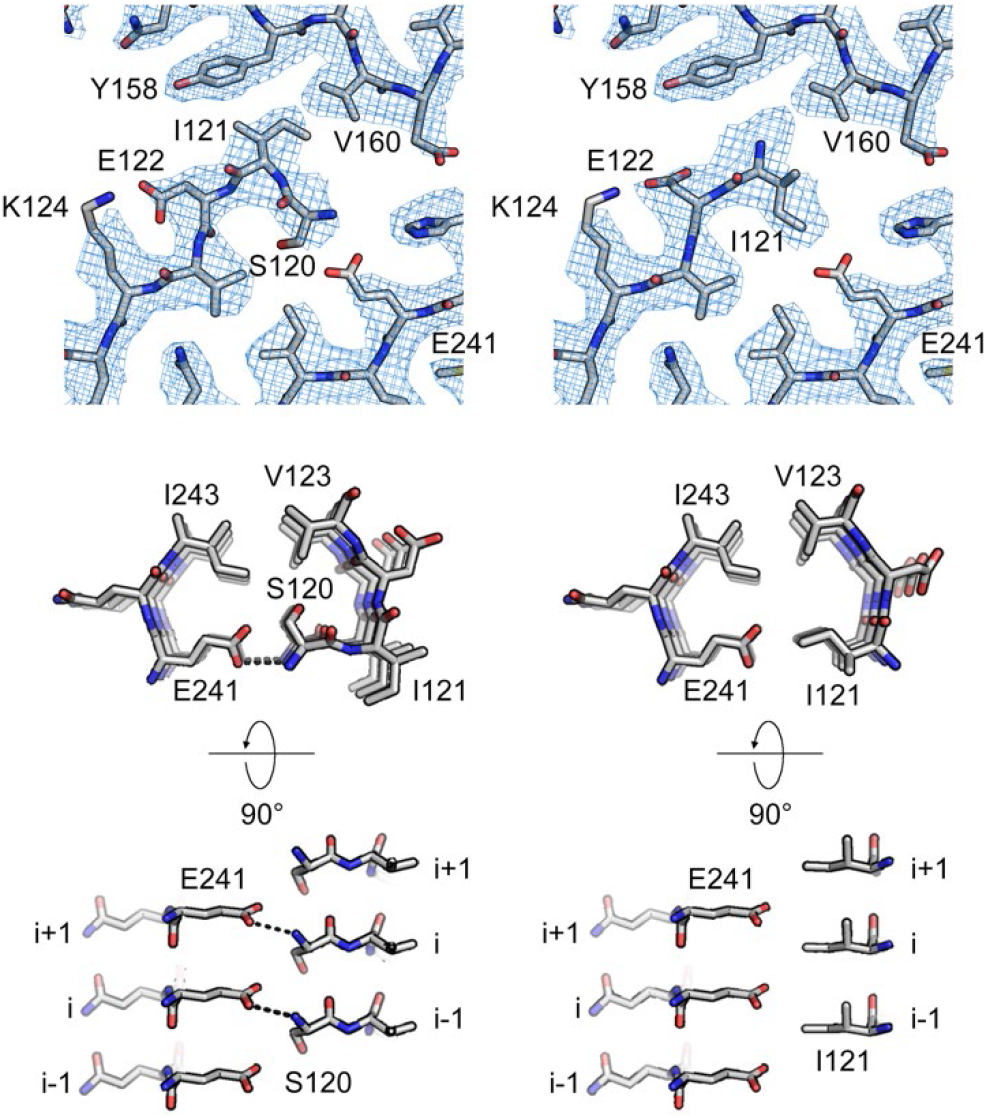
Comparison of models that start with Ser120 or Ile121 at the N-terminus. The current model starting with Ser120 (left) and an alternative model starting with Ile121 (right) are juxtaposed. We believe the N-terminus starting with Ser120 is the better model because i) Ser120 fits the density well, whereas the density is too large for only the nitrogen of Ile121; ii) Ser120 hydrogen-bonds with Glu241, whereas Ile121 does not participate in interactions that favor the fibril fold.

**Table S1.** Demographics and inclusion properties for all FTLD-TDP and non-FTLD-TDP donors.

| Original ID | Gender | Age | Braak | PathDx (type) | FHx | Brain region | Fibrils under EM | Band in western blot | Donor ID reported |
| --- | --- | --- | --- | --- | --- | --- | --- | --- | --- |
| F1 | Male | 66 | 2 | FTLD-TDP (A) | FALSE | MF | Yes | n.a. |  |
| F2 | Male | 65 | 2.5 | FTLD-TDP (A) | FALSE | MF | Yes | n.a. |  |
| F3 | Male | 60 | 3 | FTLD-TDP (A) | FALSE | MF | No | No |  |
| F4 | Female | 71 | 2 | FTLD-TDP (A) | TRUE | MF | Yes | Yes |  |
| F5 | Male | 71 | 2.5 | FTLD-TDP (A) | TRUE | MF | Yes | n.a. |  |
| F6 | Male | 61 | 1 | FTLD-TDP (A) | FALSE | MF | Yes | Yes |  |
| F7 | Male | 60 | 2 | FTLD-TDP (A) | TRUE | MF | Yes | Yes |  |
| F8 | Female | 84 | 2.5 | FTLD-TDP (A) | TRUE | MF | Yes | n.a. |  |
| F9 | Male | 76 | 1 | FTLD-TDP (A) | FALSE | MF | Yes | n.a. |  |
| F10 | Female | 78 | 1.5 | FTLD-TDP (A) | FALSE | MF | Yes | n.a. |  |
| F11 | Female | 85 | 1 | FTLD-TDP (A) | FALSE | MF | Yes | n.a. |  |
| F12 | Male | 63 | 0 | FTLD-TDP (A) | TRUE | MF | Yes | n.a. |  |
| F13 | Male | 86 | 2.5 | FTLD-TDP (A) | FALSE | MF | Yes | Yes |  |
| F14 | Male | 70 | 1 | FTLD-TDP (A) | TRUE | MF | Yes | Yes |  |
| F15 | Female | 83 | 3 | FTLD-TDP (A) | TRUE | MF | Yes | Yes |  |
| F16 | Male | 60 | 1 | FTLD-TDP (A) | TRUE | MF | Yes | Yes |  |
| F17 | Male | 86 | 0 | FTLD-TDP (A) | FALSE | MF | Yes | Yes | Donor 3 |
| F18 | Female | 90 | 2.5 | FTLD-TDP (A) | FALSE | MF | Yes | n.a. |  |
| F19 | Female | 64 | 2 | FTLD-TDP (A) | TRUE | MF | Yes | Yes |  |
| F20 | Male | 83 | 3 | FTLD-TDP (A) | FALSE | MF | Yes | Yes |  |
| F21 | Female | 71 | 0 | FTLD-TDP (C) | FALSE | MF | Yes | n.a. |  |
| F22 | Male | 70 | 0.5 | FTLD-TDP (C) | FALSE | MF | Yes | Yes |  |
| F23 | Male | 75 | 0 | FTLD-TDP (C) | FALSE | MF | Yes | Yes |  |
| F24 | Male | 74 | 2.5 | FTLD-TDP (C) | FALSE | MF | Yes | Yes |  |
| F25 | Female | 75 | 2.5 | FTLD-TDP (C) | FALSE | MF | Yes | n.a. |  |
| F26 | Male | 65 | 2 | FTLD-TDP (C) | FALSE | MF | Yes | Yes | Donor 1 |
| F27 | Male | 66 | 3 | FTLD-TDP (C) | TRUE | MF | No | No |  |
| F28 | Male | 75 | 2 | FTLD-TDP (C) | FALSE | MF | Yes | n.a. |  |
| F29 | Male | 65 | 1 | FTLD-TDP (C) | FALSE | MF | Yes | Yes |  |
| F30 | Male | 78 | 1 | FTLD-TDP (C) | TRUE | MF | Yes | Yes |  |
| F31 | Female | 69 | 3 | FTLD-TDP (C) | FALSE | MF | Yes | Yes |  |
| F32 | Male | 66 | 0 | FTLD-TDP (C) | FALSE | MF | Yes | Yes |  |
| F33 | Female | 69 | 2 | FTLD-TDP (C) | TRUE | MF | Yes | n.a. |  |
| F34 | Female | 82 | 3 | FTLD-TDP (C) | TRUE | MF | Yes | Yes |  |
| F35 | Male | 70 | 2.5 | FTLD-TDP (B) | TRUE | MF | Yes | No |  |
| F36 | Female | 76 | 3 | FTLD-TDP (B) | FALSE | MF | Yes | Yes | Donor 2 |
| F37 | Female | 62 | 0 | FTLD-TDP (B) | FALSE | MF | Yes | n.a. |  |
| F38 | Female | 62 | 1 | FTLD-TDP (B) | TRUE | MF | Yes | n.a. |  |
| F39 | Male | 77 | 2 | FTLD-TDP (B) | TRUE | MF | Yes | Yes |  |
| F40 | Female | 64 | 3 | FTLD-TDP (D) | TRUE | MF | Yes | Yes | Donor 4 |
| N1 | Male | 69 | 2 | Normal | No | MF | No | No |  |
| N2 | Male | 62 | 2 | VaD | Yes | MF | No | No |  |
| N3 | Male | 79 | 1 | VaD | No | MF | No | No |  |
| N4 | Female | 87 | 2 | VaD | No | MF | No | No |  |
| N5 | Male | 59 | 0 | Normal | No | MF | No | No | Donor 5 |
| N6 | Male | 53 | 1 | PART | No | MF | No | No |  |
| N7 | Female | 74 | 2 | PART | No | MF | No | No |  |
| N8 | Male | 74 | 3 | PART | No | MF | No | No |  |

PathDx: pathological diagnosis; FHx: family history; VaD: vascular dementia; PART: primary age-related tauopathy; MF: medial frontal gyrus; Band in western blot: detection of the ~35 kDa TMEM106B antibody-positive band in the sarkosyl-insoluble fraction; n.a.: not available. All donors were numbered by original ID (F1-40 for FTLD-TDP cases and N1-8 for non-FTLD-TDP cases) during initial fibril screen. F26, F36, F17, F64 and N5 were selected for further study and re-numbered as donor 1-5 in the main text.

**Table S2.** Cryo-EM data collection and processing statistics of FTLD-TDP donors 2-4.

|  | Donor 2 |  |  | Donor 3 |  |  | Donor 4 |  |  |
| --- | --- | --- | --- | --- | --- | --- | --- | --- | --- |
|  | PM1 | PM2 | PM3 | PM1 | PM2 | PM3 | PM1 | PM2 | PM3 |
| Magnification | ×81,000 | ×81,000 | ×81,000 | ×81,000 | ×81,000 | ×81,000 | ×81,000 | ×81,000 | ×81,000 |
| Voltage (kV) | 300 | 300 | 300 | 300 | 300 | 300 | 300 | 300 | 300 |
| Electron exposure (e<sup>-</sup>/Å<sup>2</sup>) | 37 | 37 | 37 | 36 | 36 | 36 | 36 | 36 | 36 |
| Defocus range (μm) | 1.0-5.1 | 1.0-5.1 | 1.0-5.1 | 0.5-5.1 | 0.5-5.1 | 0.5-5.1 | 0.8-4.9 | 0.8-4.9 | 0.8-4.9 |
| Pixel size (Å) | 1.1 | 1.1 | 1.1 | 1.1 | 1.1 | 1.1 | 1.1 | 1.1 | 1.1 |
| Symmetry imposed | C<sub>1</sub> | C<sub>2</sub> | C<sub>1</sub> | C<sub>1</sub> | C<sub>2</sub> | C<sub>1</sub> | C<sub>1</sub> | C<sub>2</sub> | C<sub>1</sub> |
| Helical rise (Å) | 4.896 | 4.9 | 2.446 | 4.897 | 4.906 | 2.452 | 4.897 | 4.904 | 2.4281 |
| Helical twist (°) | -0.413 | 179.619 | 179.807 | -0.413 | 179.6 | 179.8 | -0.413 | 179.605 | 179.81 |
| Initial particle images (no.) | 438,393 | 438,393 | 438,393 | 419,107 | 419,107 | 419,107 | 228,432 | 228,432 | 228,432 |
| Final particle images (no.) | 38,894 | 24,420 | 5,317 | 26,533 | 15,587 | 5,335 | 14,407 | 13,482 | 5,798 |
| Map resolution (Å) | 3.8 | 4.0 | 4.6 | 3.6 | 4.1 | 5.3 | 3.5 | 3.6 | 5.0 |
| FSC threshold | 0.143 | 0.143 | 0.143 | 0.143 | 0.143 | 0.143 | 0.143 | 0.143 | 0.143 |
| Map resolution range (Å) | 200-3.8 | 200-4.0 | 200-4.6 | 200-3.6 | 200-4.1 | 200-5.3 | 200-3.5 | 200-3.6 | 200-5.0 |

**Table S3.**
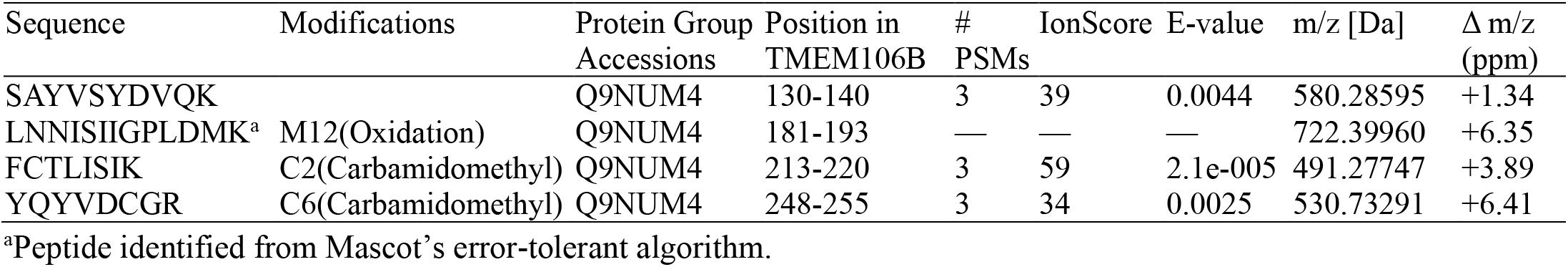
TMEM106B peptides identified by mass spectrometry.

**Table S4.** BLAST search results.

| Description | Scientific Name | Max Score | Total Score | Query Cover | E-value | Per. Ident | Accession Length | Accession |
| --- | --- | --- | --- | --- | --- | --- | --- | --- |
| TMEM106B protein | Homo sapiens | 39.3 | 39.3 | 93% | 0.003 | 16.00% | 274 | AAH39741.1 |
| transmembrane protein 106B | Homo sapiens | 39.3 | 39.3 | 93% | 0.003 | 16.00% | 274 | NP_001127704.1 |
| transmembrane protein 106B, isoform CRA_b | Homo sapiens | 39.3 | 39.3 | 92% | 0.003 | 16.13% | 285 | EAW93639.1 |
| TMEM106B protein | Homo sapiens | 38.9 | 38.9 | 93% | 0.004 | 16.00% | 314 | AAH28108.1 |
| hypothetical protein FLJ11273 variant | Homo sapiens | 37.0 | 37.0 | 92% | 0.019 | 16.13% | 274 | BAD96983.1 |
| nuclear receptor subfamily 2 group E member 1 isoform b | Homo sapiens | 25.4 | 25.4 | 50% | 224 | 32.89% | 385 | NP_003260.1 |

